# Unprecedented biomass and fatty acid production by the newly discovered cyanobacterium *Synechococcus* sp. PCC 11901

**DOI:** 10.1101/684944

**Authors:** Artur Włodarczyk, Tiago Toscano Selão, Birgitta Norling, Peter J. Nixon

## Abstract

Cyanobacteria, which use solar energy to convert carbon dioxide into biomass, are potential solar biorefineries for the sustainable production of chemicals and biofuels. However, yields obtained with current strains are still uncompetitive compared to existing heterotrophic production systems. Here we report the discovery and characterization of a new cyanobacterial strain, *Synechococcus* sp. PCC 11901, with potential for green biotechnology. It is transformable, has a short doubling time of ≈2 hours, grows at high light intensities and a wide range of salinities and accumulates up to 18.3 g dry cell weight/L of biomass – 2-3 times more than previously described for cyanobacteria - when grown in a modified medium containing elevated nitrate, phosphate and iron. As a proof of principle, PCC 11901 engineered to produce free fatty acids did so at unprecedented levels for cyanobacteria, with final yields reaching over 6 mM (1.5 g/L), comparable to those achieved by heterotrophic organisms.

## 1. Introduction

The current production of commodity chemicals from fossil fuels results in the release of greenhouse gases, such as CO_2_, into the atmosphere. Given concerns over the possible link between greenhouse gases and climate change and the limited abundance of cheap fossil fuels^1^, alternative sustainable approaches need to be developed to produce carbon-based chemicals on an industrial scale. Yeast and bacteria are widely used biotechnology production platforms. However, their growth relies on the addition of carbohydrates to the growth medium leading to a ’food vs. fuel’ dilemma, which will likely drive the price of most carbon feedstocks up, fueled by the negative impact of global warming on food crops^2^. Cyanobacteria have the potential to provide a completely sustainable solution^3, 4^. These evolutionary ancestors of algal and plant chloroplasts are gram-negative prokaryotic oxyphotoautotrophs, able to convert carbon dioxide and inorganic sources of nitrogen, phosphorous and microelements into biomass^5^.

Among cyanobacterial strains, the marine strain *Synechococcus* sp. PCC 7002^6^ and several freshwater strains, namely *Synechocystis* sp. PCC 6803^7^, *Synechococcus elongatus* PCC 7942^8^ and more recently *Synechococcus elongatus* UTEX 2973^9^, have become model organisms for both basic photosynthesis research as well as the photoautotrophic production of different chemicals such as bioplastics^10^, biofuels (ethanol^11^ and free fatty acids^12, 13^) and specialized compounds like terpenoids^14, 15^. However, the average yields are often low compared to heterotrophic microbes, partly due to relatively slow growth and low biomass accumulation.

In this work, we report the isolation and detailed characterization of the novel cyanobacterial strain *Synechococcus* sp. PCC 11901 (hereafter PCC 11901) and demonstrate its potential for cyanobacterial biotechnology. PCC 11901 tolerates temperatures up to a maximum of 43 °C, high light irradiances and salinities, with average doubling times ranging from 2-3 hours in 1% CO_2_/air. We have also conducted a parametric analysis to devise a new culture medium, termed MAD medium, which enables cultures of PCC 11901 to produce the highest value of dry cell biomass of any cyanobacterial strain described so far. Lastly, we tested the ability of PCC 11901 to efficiently convert CO_2_ into free fatty acids (FFA), a class of commodity chemical used by a range of industries^16^. Although its current production from palm oil can be considered as renewable^17^, growing world demand and extensive farming of palm oil trees in South East Asia are causing irreversible deforestation of primordial rainforests and degradation of the natural habitats in producing countries^18^. Yields of FFA obtained with PCC 11901 reached 6.16 mM (≈1.54 g/L) after 7 days of cultivation, in the same range of productivity attained by heterotrophic hosts and 2.4^13^ to 11.8-fold^12^ greater than yields previously achieved with cyanobacteria.

## 2. Results

### 2.1. Strain isolation, identification and characterization

As our primary goal was to isolate a fast-growing, preferably marine cyanobacterial strain that would not compete for freshwater resources and could tolerate a wide range of abiotic stresses, we collected samples at a local floating fish farm (an environment rich in waste nitrogen and phosphorous compounds), located in the Johor river estuary in Singapore. Enrichment of water samples and consecutive re-streaking on solid medium (see Materials and Methods) resulted in the isolation of a fast-growing cyanobacterial colony. Inoculation into liquid AD7 medium lacking an organic carbon source, which is used widely to grow marine cyanobacteria^19, 20^ led to rapid growth, but only after a long lag phase (≈24 hours), suggesting nutrient limitation. As most oceanic phytoplankton require cobalamin (vitamin B12) for growth, and its concentrations in the seawater and oceans vary from 0 to 3 pM^21, 22^, we supplemented cultures of the isolated xenic strain with 3 pM cobalamin, which alleviated the lag phase. Upon dilution plating of the culture on solid medium with cobalamin, small transparent bacterial colonies were found in the vicinity of the cyanobacterial colonies and both strains were further purified by streaking on a 1:1 mixture of AD7 and LB medium. 16S rRNA sequencing coupled to phylogenetic analyses revealed that the axenic cyanobacterial strain was a member of the *Synechococcales* group (now deposited in the Pasteur Culture Collection as *Synechococcus* sp. PCC 11901) and that the closest phylogenetic relatives to the companion heterotrophic bacterial strain belong to the marine *Thalassococcus* genus^23^. Both the axenic and xenic strains of PCC 11901 grew equally well in the presence of cobalamin, reaching an OD_730_ ≈23 after 72 hours (Fig. 1a). In contrast, growth of the axenic strain was severely inhibited in the absence of added cobalamin with the observed residual growth probably reflecting the retention of intracellular cobalamin or methionine reserves^24^ in the inoculum (Fig. 1a). These results imply that PCC 11901 is auxotrophic, and that its growth depends on the availability of cobalamin in the growth medium.

**Fig. 1.**
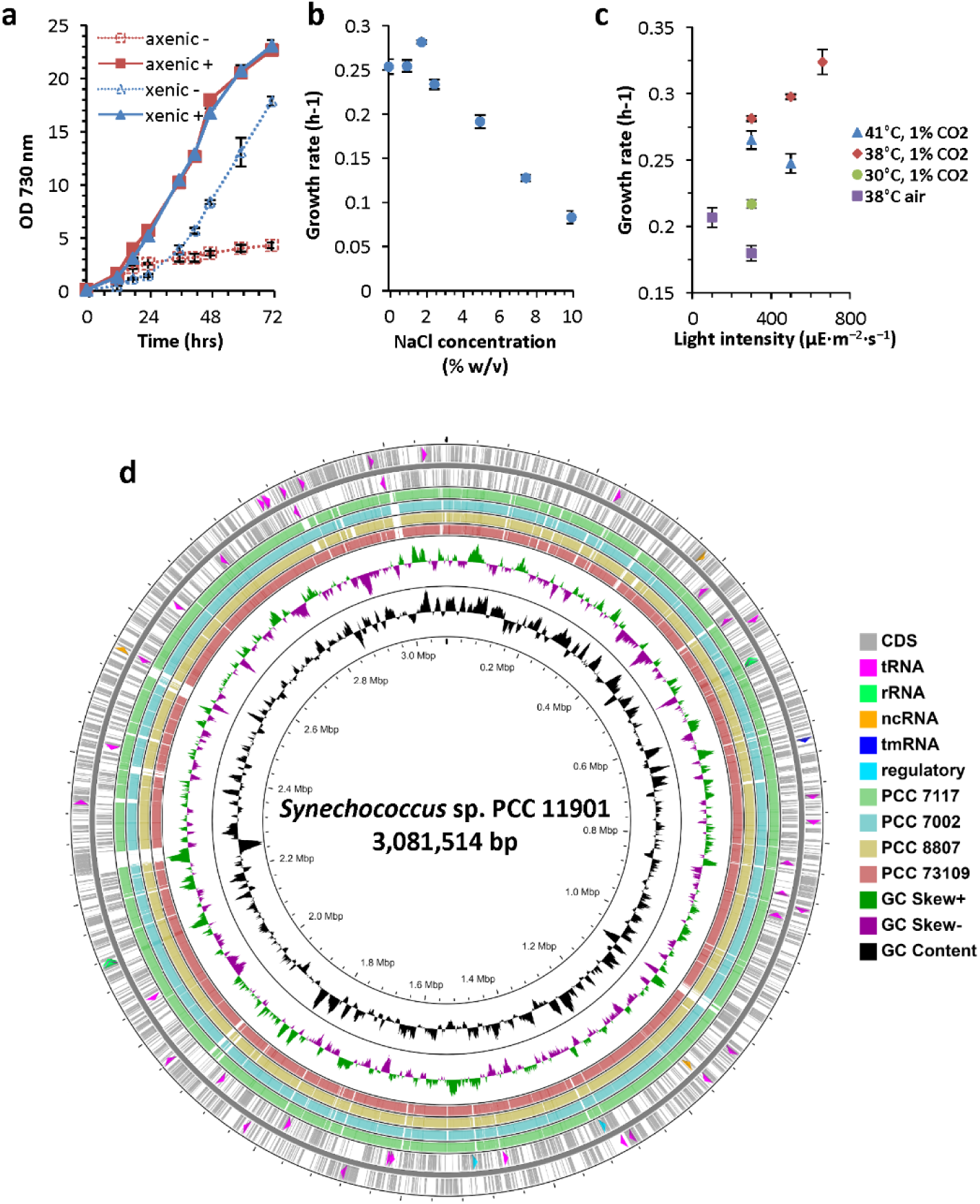
Growth and genome analysis of *Synechococcus* sp. PCC 11901 strain. **(a)** Axenic and xenic (contaminated with *Thalassococcus sp.*) strains grown in AD7 medium with (+) and without (-) addition of cobalamin (vitamin B_12_). **(b)** Growth rate of axenic strain grown in AD7 medium with different concentrations of sodium chloride (0-10%). For growth curves see **Fig. S1**. Growth conditions for **(a)** and **(b)**: 38 °C, 225 rpm, 300 µmol photons·m^-2^·s^-1^ light intensity (RGB LED 4:2:1 ratio) and 1% CO_2_. **(c)** Growth rates of the axenic strain cultured under different light, temperature and CO_2_ conditions. For all growth curves and doubling times see **Fig. S2** and **Table S1**. Data points with error bars represent mean of n=3 biological replicates ± standard deviation. **(d)** Circular diagram of the PCC 11901 genome sequence combined with the BLAST scores of four related cyanobacterial strains compared to PCC 11901 genome. Tracks: CDS (grey), tRNA (pink), rRNA (green), ncRNA (orange), tmRNA (dark blue), regulatory elements (turquoise), GC Skew (above average: green, below average: purple), GC content (black). BLAST scores: PCC 7117 (green), PCC 7002 (light blue), PCC 8807 (tan) and PCC 73109 (pink) compared to the PCC 11901 genome. Image created using CGView Server^BETA^ online tool^28^.

Although we did not sequence the genome of the isolated *Thalassococcus* strain, another *Thalassococcus* sp. strain (SH-1) possesses cobalamin biosynthesis genes (GenBank accession number CP027665.1). Given the previous evidence for the mutualistic interaction between cobalamin-dependent microalgae and bacteria^22^, it is very likely that the *Thalassococcus* contaminant in the xenic culture could serve as a natural symbiotic partner for the PCC 11901 strain providing it with cobalamin, while consuming nutrients excreted by the cyanobacterium.

As in the case of PCC 7002^25^, the PCC 11901 strain can grow mixotrophically in the presence of glycerol and photoheterotrophically in the presence of glycerol and the herbicide DCMU, which inhibits photosynthetic electron flow, although growth is impaired **(Fig. S3c)**. Interestingly, this strain could not utilize glucose for growth, as very little or no growth was observed after one week of incubation with glucose and DCMU **(Fig. S3d).**

Analysis of cell morphology revealed that PCC 11901, though unicellular, could form short filaments of 2-6 cells, depending on growth phase (**Fig. S11**). Individual cells are elongated with sizes ranging from 1.5 to 3.5 µm in length and 1-1.5 µm in width. On average (n=45), cells contain 4 to 6 concentric layers of thylakoids around the cytoplasm with visible convergence zones on the cell periphery (Fig. 2a-c). In negatively stained cells, long (1-1.5 µm) fibres of pili-like structures, similar to those seen previously in other cyanobacteria^26, 27^, were observed extending from the outer cell wall membrane (Fig. 2d-e).

**Fig. 2.**
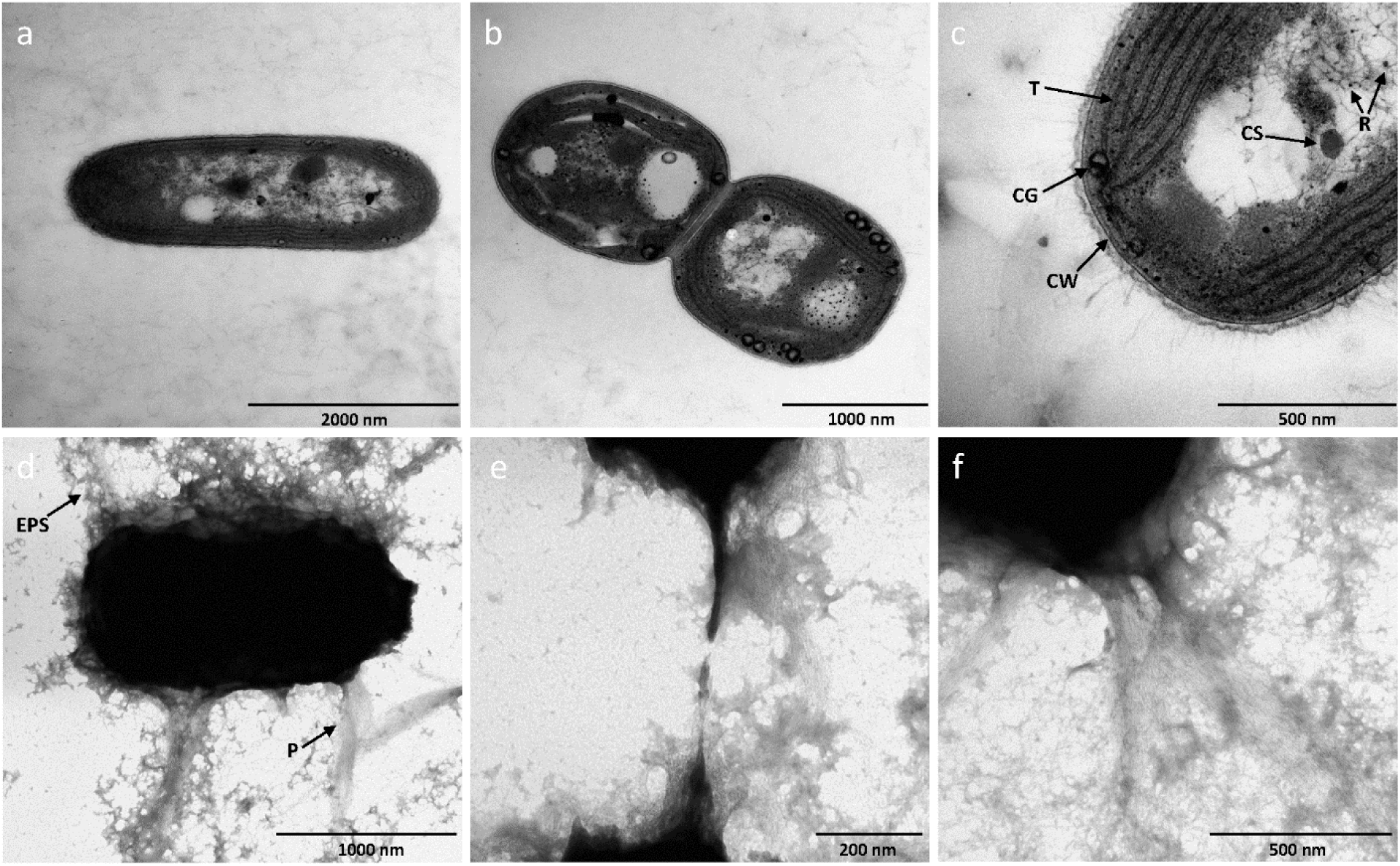
Transmission electron micrographs. **(a-c)** Cross section of the cells. **(d-f)** Negatively stained cells. Abbreviations: CW – cell wall, T – thylakoids, CS – carboxysome, R – ribosome, CG – cyanophycin granule, EPS – exopolysaccharide-like structures, P – pili-like structures.

### 2.2. Sequencing, phylogenetic assignment and genome analysis

The genome of the PCC 11901 strain was sequenced using both Illumina and PacBio next generation sequencing techniques, assembled as described in Materials and Methods and deposited in GenBank under accession number CP040360.1. Two sequences (one chromosome and one plasmid) were fully covered and circularized, and four shorter sequences (most likely plasmids) with sizes ranging from 10 to 98 kbp could not be closed. The complete genome size is 3,081,514 bp, similar in size to other phylogenetically related cyanobacterial strains, with an average GC content of 49.5%. The chromosome contains 42 tRNA, 6 rRNA, 4 ncRNA and 2,943 gene sequences (3,316 including plasmids). Distribution of all genetic elements on the genome is shown on 2 peripheral circles corresponding to strand directions in Fig. 1d.

The average nucleotide identity (ANI) analysis of the chromosome revealed 97.48% identity to *Synechococcus* sp. PCC 7117 strain, 96.99% to both PCC 73109 and 8807, 96.76% to the commonly used PCC 7002 strain and 90.03% to PCC 7003. Complete genome sequences of the above mentioned strains were compared to the PCC 11901 strain genome using BLAST analysis in the CGView Server^28^ tool. Rather unusually, there are several major insertions in the genome of the PCC 11901, depicted by white gaps in the BLAST search scores shown in Fig. 1d, the biggest of approximately 24 kbp, which are not found in other cyanobacterial genomes. Analysis of these fragments showed similarity to genes from other cyanobacteria outside the *Synechococcales* taxonomy order group, with a strong prevalence of genes encoding predicted glycosyltransferase genes (14 genes), ABC transporter components, transposases, toxin-antitoxin system components and an alcohol dehydrogenase (**Table S5**).

In one of the endogenous plasmids (GenBank accession number: CP040361.1) we identified a gene encoding a 5-methyltetrahydropteroyltriglutamate-homocysteine S-methyltransferase homologue (MetE, GenPept: QCS51047.1) involved in cobalamin-independent methionine synthesis. Although most cyanobacteria are capable of synthesizing cobalamin or pseudocobalamin required by the cobalamin-dependent methionine synthase (MetH), some strains, such as PCC 7002, lack MetE, and are therefore strict cobalamin auxotrophs^24, 29^. Surprisingly, despite having both variants of methionine synthase in the genome, PCC 11901 is dependent on added cobalamin for growth (Fig. 1a), suggesting potential mutations in either the *metE* gene itself or in a corresponding cobalamin riboswitch leading to either a loss of activity or expression of this gene. Interestingly, two genes encoding enzymes in the cobalamin biosynthetic pathway *cobA* (encoding uroporphyrinogen-III C-methyltransferase, GenPept: QCS48576.1) and *cobQ* (encoding cobyric acid synthase, GenPept: QCS50783.1) can be found in the chromosome, suggesting that they could have been either acquired from other strains by horizontal gene transfer or the cobalamin biosynthesis pathway could have been lost during evolution.

### 2.3. Growth performance under different salinity, temperature and light conditions

To test the tolerance of PCC 11901 to high salt concentrations, growth was assessed in the presence of a wide range of NaCl concentrations, from 0 to 10% (w/v), where sea water is equivalent to ≈3.5 % NaCl (Fig. 1b). PCC 11901 growth rates did not differ significantly in the 0-2.5% NaCl concentration range, with a maximum of 0.28 h^-1^ at 1.8% NaCl. Higher NaCl concentrations had a noticeably negative impact on growth rates, although cultures grown in the presence of 7.5% NaCl were still able to reach an OD_730_ of 17.4 after 3 days of cultivation and even in the presence of 10% NaCl were able to grow to a final OD_730_ = 6.3 with an average doubling time of 12 hours, a similar growth rate to that of the halotolerant microalga *Dunaliella*^30^.

Different combinations of temperature, light intensity and CO_2_ conditions were tested in order to investigate their effect on growth (Fig. 1c). As a reference, PCC 11901 was grown alongside two other fast-growing and high-temperature tolerant strains – UTEX 2973 and PCC 7002 (**Fig. S2**). All strains grown in atmospheric CO_2_ conditions exhibited similar doubling times with the shortest of 3.35 ± 0.12 hours at low light (100 μmol photons·m^−2^·s^−1^) observed for PCC 11901 (**Table S1**). At higher light the UTEX 2973 strain grew faster in atmospheric CO_2_ conditions, though this result could be partially influenced by the presence of both sodium carbonate and citric acid in BG-11 medium (used to grow freshwater strains) and the absence of such carbon sources in AD7 medium (used to grow marine strains). PCC 11901 was found to grow the fastest at 38 °C and 1% CO_2_, with a shortest doubling time of 2.14 ± 0.06 hours observed at a light intensity of 660 μmol photons·m^−2^·s^−1^. The highest permissible growth temperature for PCC 11901 was 43 °C. Under our growth conditions, UTEX 2973 was the fastest growing cyanobacterium, with a shortest doubling time of 1.93 ± 0.04 hours at 41 °C and 1% CO_2_. However, this advantage over PCC 11901 and 7002 strains becomes less apparent when grown for longer periods. After 24 hours of cultivation UTEX 2973 reached lower OD_730_ values than the other two tested strains and was unable to exceed OD_730_ ≈ 10 after 4 days (**Fig. S4**), an effect that was also previously observed^31^. Regardless of light conditions, PCC 11901 accumulated more biomass after 4 days of growth (4.9 g/L) than PCC 7002 and UTEX 2973 (3.7 and 2.5 g/L respectively). However, all strains accumulated less biomass under higher light, especially PCC 7002 (**Fig. S4**).

### 2.4. Transformability and availability of molecular tools for protein expression

For a new strain to be useful for synthetic biology and metabolic engineering applications, it must be amenable to transformation and have molecular tools for controlled protein expression. Considering that PCC 11901 is a close relative of PCC 7002, we tested several molecular tools already established for PCC 7002. In one construct (Fig. 3a) the *yfp* gene was placed under control of the constitutive P_cpt_ promoter (a truncated *cpcB* promoter from the PCC 6803 strain functional in PCC 7002)^32^ and inserted between flanking regions of the *acsA* (acetyl-CoA synthetase) gene. As previously reported, targeting of genes to the *acsA* locus in PCC 7002 allows for the generation of markerless mutants using acrylic acid for counterselection^33^. In the second construct, the *yfp* gene and a spectinomycin-resistance cassette were cloned between flanking regions of the *psbA2* (encoding the D1 subunit of photosystem II) gene and YFP expression was controlled by the synthetic IPTG-inducible P_clac143_ promoter^32^. PCC 11901 was then transformed by natural transformation (with transformation efficiency similar to PCC 7002) and positive clones were confirmed by colony PCR **(Fig. S10a-b)**. Fluorescence microscopy confirmed successful YFP expression in both constructs (Fig. 3b), with expression in the Δ*psbA2*::P_clac143_-YFP strain detectable only after induction with IPTG. Though the targeted genome loci are different for these two constructs, the P_cpt_ promoter appeared to be stronger than the induced P_clac143_ promoter, as the relative fluorescence unit (RFU) per OD_730_ ratio was higher (Fig. 3c). YFP under control of the P_clac143_ promoter was expressed upon induction at 45-fold greater levels than the non-induced control (Fig. 3c).

**Fig. 3.**
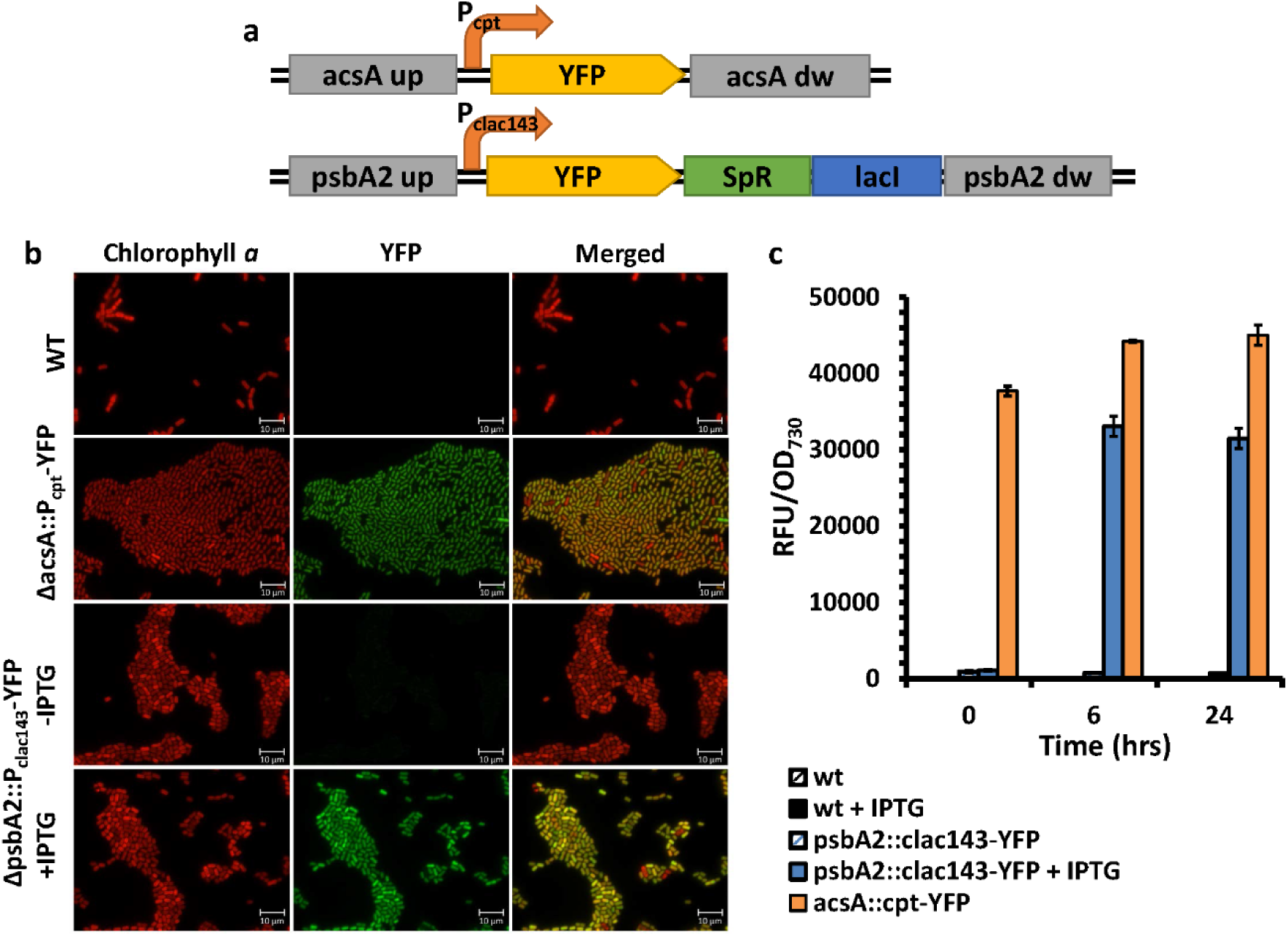
Usability of different molecular tools for the expression of heterologous proteins in *Synechococcus* sp. PCC 11901. **(a)** Schematic illustration of constructs used for natural transformation and homologous recombination in the chromosome of 11901 strain. In the pSW036 vector YFP gene was inserted between 700 bp flanking regions of the *acsA* gene and expression was controlled by the synthetic and constitutive P_cpt_ promoter. In the pSW039 vector YFP gene following synthetic, inducible P_clac143_ promoter together with *lacI* regulator and spectinomycin-resistant cassette were inserted between 700 bp flanking regions of the *psbA2* gene. **(b)** Fluorescence microscopy images of cells transformed with YFP constructs. As a control, chlorophyll *a* autofluorescence of WT, Δ*acsA*::P_cpt_-YFP and Δ*psbA2*::P_clac143_-YFP is shown on the left side of the panel, and YFP fluorescence is shown in the middle panel. Δ*psbA2*::P_clac143_-YFP strain was incubated for 24 hours with/without addition of 1 mM IPTG prior to imaging. **(c)** Strength and inducibility comparison of P_cpt_ and P_clac143_ promoters. Relative YFP fluorescence in relation to OD_730_ was measured for cultures at OD ≈1 (T=0), after 6 and 24 hours. The bars represent mean of n=3 biological replicates ± standard deviation.

### 2.5. Identification of nutrient constraints

Modifying the composition of the growth medium is a common approach to improve biomass and secondary metabolite production using heterotrophic microorganisms^34^. In contrast, the media routinely used to cultivate cyanobacteria were formulated over 40 years ago and are not optimised for high biomass production^29^. Indeed modification of the basic AD7 medium by Clark et al.^31^ was shown to increase biomass production of PCC 7002 by approximately 2-fold with further improvements theoretically possible. Other work has also described an enriched BG-11 medium for high-biomass and cyanophycin production by the freshwater strain PCC 6803^35^.

Inspired by these reports we adopted a systematic approach to improve growth of PCC 11901 by independently changing the levels of nitrate, phosphate and iron in AD7 medium (Fig. 4). These experiments led to the development of a **<u>M</u>**odified **<u>AD</u>**7 medium (hereafter named MAD) containing 96 mM NaNO_3_, 240 µM FeCl_3_ and 1.2 mM phosphate. In contrast the recently described medium A described by Clark et al. (“modified medium A” or “MMA” medium^31^) contains 122 mM NaNO_3_, 1.1 mM ammonium iron citrate and 31 mM of phosphate (by adding 5.2 mM in a fed batch scheme). We found that phosphate at levels greater than 0.8-1.2 mM had an adverse effect on growth of PCC 11901 (Fig. 4d **& S5)**. The highest biomass accumulation was achieved after 8 days of cultivation in MAD medium (12.3 gDW/L) while in the case of MMA medium containing 5.2 mM phosphate, 10.3 gDW/L was obtained in 8 days and 10.3 g/L in 10 days when supplemented with 31 mM phosphate (**Fig. S5**). This difference in biomass could reflect precipitation of iron, magnesium, calcium and other micronutrients as phosphate salts in MMA medium, thus lowering their bioavailability in basic pH.

**Fig. 4.**
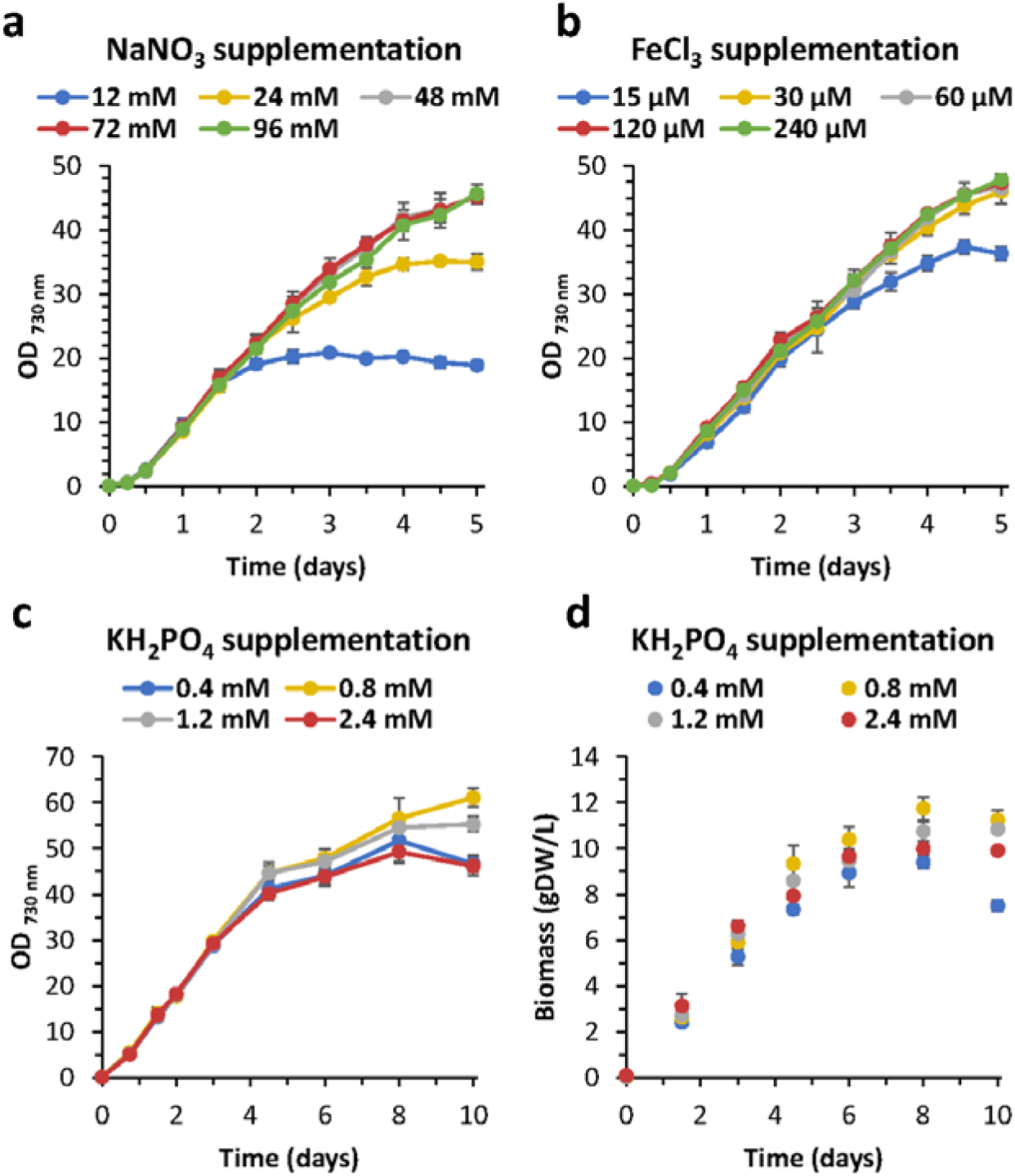
Effect of medium on growth of *Synechococcus* sp. PCC 11901. Analysis of the effect of medium supplementation with increasing concentrations of sodium nitrate, iron (III) chloride and potassium dihydrogen phosphate on growth of PCC 11901. Growth conditions: 38 °C, 225 rpm, 300 µmol photons·m^-2^·s^-1^ light intensity (RGB LED 4:2:1 ratio) and 1% CO_2_. **(a)** AD7 medium with double the concentration of FeCl_3_ (30 µM) was supplemented with a wide spectrum of NaNO_3_ concentrations. **(b)** AD7 medium enriched with 96 mM NaNO_3_ was supplemented with different concentrations of FeCl_3_. **(c,d)** Modified AD7 medium with 96 mM NaNO_3_ and 240 µM FeCl_3_ was supplemented with increasing concentrations of KH_2_PO_4_ and tested for growth and biomass accumulation over 10 days. Data points with error bars represent mean of n=3 biological replicates ± standard deviation.

### 2.6. High biomass production in MAD medium

To evaluate the general use of modified media for growing cyanobacteria (MAD for marine strains and a 5x concentrated BG-11 medium (5xBG) for freshwater strains), other commonly used strains – marine PCC 7002 and three freshwater strains PCC 7942, UTEX 2973 and PCC 6803 – were grown alongside PCC 11901 at 30 °C (the optimal temperatures for PCC 7942 and PCC 6803) and with 1% CO_2_/air. PCC 11901 clearly outperformed all the other model cyanobacteria, growing to an OD_730_ = 101 and accumulating a maximum of 18.3 gDW/L of biomass, almost twice the biomass of PCC 7002 (9.3 gDW/L) under the same conditions (Fig. 5a-b). Among the freshwater cyanobacteria, PCC 6803 accumulated the highest biomass (6.9 gDW/L), followed by PCC 7942 and UTEX 2973 – both around 6.5 gDW/L. In regard to culture fitness, loss of the light-harvesting phycobilisome complex, which is symptomatic of nitrogen stress, was already apparent in the case of UTEX 2973 after 3-4 days of cultivation but less apparent in the PCC 7942 and 11901 strains, even after 10 days of cultivation (Fig. 5c-d).

**Fig. 5.**
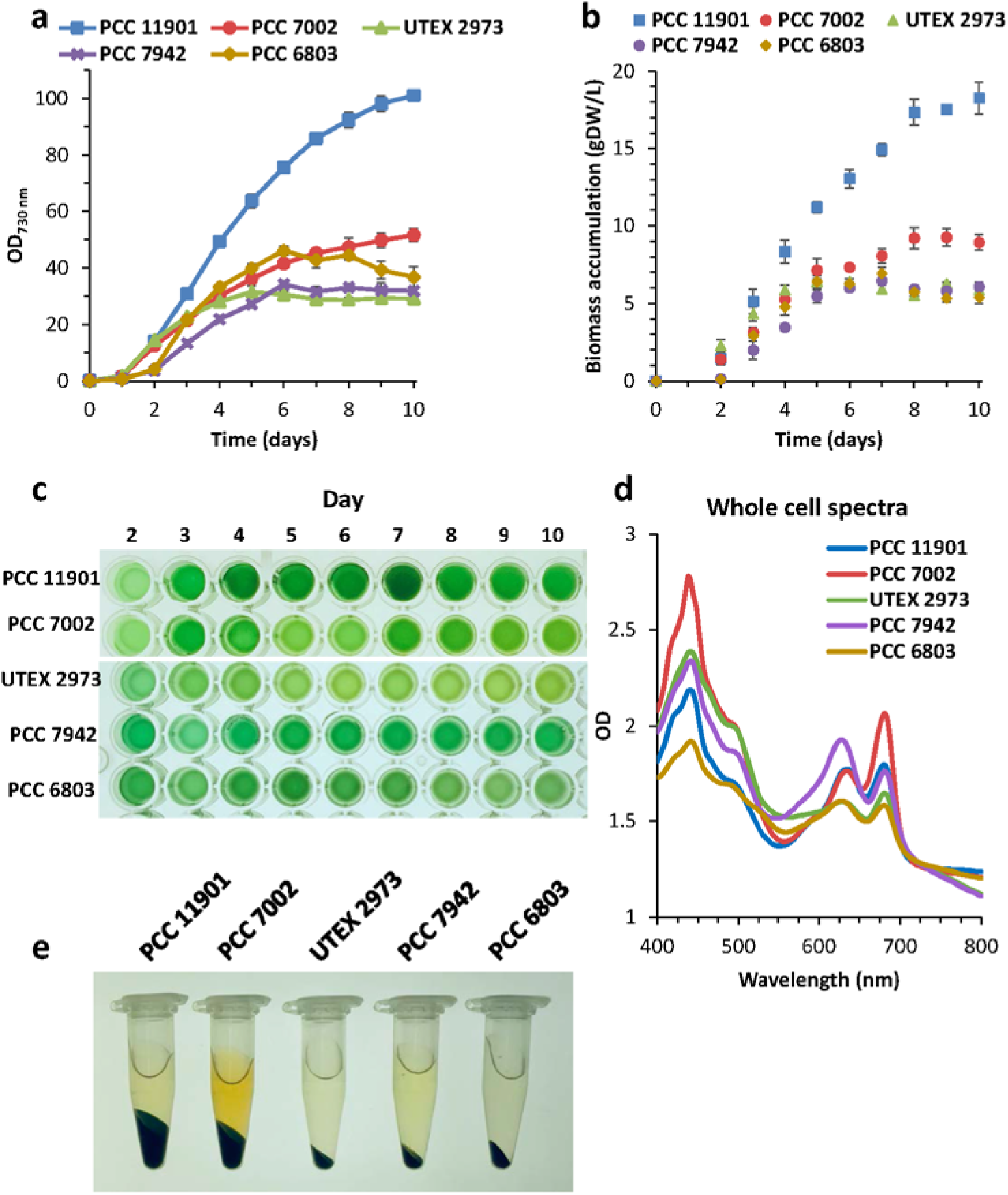
Comparison of growth performance and biomass accumulation of commonly used cyanobacteria cultured in the optimized medium. All strains were grown in triplicates at 30 °C, 200 rpm shaking under continuous illumination using RGB LED 1:1:1 ratio. *Synechcoccus* sp. PCC 11901 and 7002 were grown in the MAD medium at initially 150 µmol photons·m^-2^·s^-1^, which was then increased to 750 µmol photons·m^-2^·s^-1^ after 1 day. *Synechococcus elongatus* UTEX 2973 was grown in 5xBG medium under the same conditions. *Synechococcus elongatus* PCC 7942 and *Synechocystis* sp. PCC 6803 were also grown in 5xBG medium, with light intensity gradually increased from 75 to 150 µmol photons·m^-2^·s^-1^ (day 1) and eventually set to 750 µmol photons·m^-2^·s^-1^ (day 2). **(a, b)** Growth curve and biomass accumulation comparison of all tested cyanobacterial strains. **(c)** Culture samples over 10 days of cultivation. **(d)** Whole cell spectra of cultures harvested after 10 days of cultivation and diluted to the same OD_730_. **(e)** Comparison of the cell pellet size from 1 mL of the cultures harvested after 10 days of cultivation. Data points with error bars represent mean of n=3 biological replicates ± standard deviation.

Despite numerous efforts, we were unable to grow any of the freshwater cyanobacteria used in this study to biomass levels exceeding 7 gDW/L. Medium optimization appeared to be more challenging, possibly due to a lower tolerance to higher ionic strength in the media by these strains. We tested five different media formulations for UTEX 2973, PCC 7942 and PCC 6803 (**Fig. S7**, **Table S2**). Supplementing BG-11 medium with more nitrate, phosphate, ammonium iron (III) citrate and magnesium sulphate was met with limited success, as, in spite of a fast intial growth, cultures declined dramatically after a few days **(Fig. S7a-c)**. Cultures grown in modified MAD medium lacking sodium chloride did not perform well either **(Fig. S7d)**. In our hands, 5xBG was the most successful formulation (**see** Fig. 5 **and S7e-f**) and to our surprise supplementing this medium with phosphate and nitrate concentrations to the levels in MAD medium (modified 5xBG, 5xBGM) led to complete bleaching of the PCC 6803 strain after 10 days **(Fig. S8a-b)**. Since the 5xBGM medium contains more nitrate and phophate than 5xBG, it is very unlikely that the cultures bleached as a result of the nutrient starvation. Lippi et al.^35^ have previously shown that supplementation of BG-11 medium with 65 mM nitrate and 10 mM phosphate allows PCC 6803 to grow to high cell densities (OD=40) though, again, not surpassing the density here observed with 5xBG. Therefore, it is possible that these freshwater strains may have some limitations related to low tolerance to high concentration of inorganic salts or a regulatory mechanism (e.g. quorum sensing) preventing them from growing further.

### 2.7. Photoautotrophic production of free fatty acids

Having demonstrated that the PCC 11901 strain is amenable to genetic manipulation, a proof of concept demonstration of the strain’s biotechnological potential was designed. For that purpose, we chose to modify PCC 11901 for photoautotrophic production of FFA, which can potentially be used as renewable industrial feedstocks^16^. We therefore inserted a truncated version of the *E. coli* thioesterase (‘*tesA*) gene^12^, codon-optimized for PCC 7002, under control of the inducible P_clac143_ promoter^32^, in the genomes of both PCC 11901 and PCC 7002, by simultaneously knocking-out of the long-chain-fatty-acid-CoA ligase (*fadD*), to generate 11901 Δ*fadD*::*tesA* and 7002 Δ*fadD*::*tesA* strains (Fig. 6a). Knockout strains of the *fadD* gene alone (Δ*fadD*) and WT were used as production controls. In order to check if the MAD medium would also allow an improvement of the production yields, all strains were grown side-by-side using either regular AD7 or MAD medium.

**Fig. 6.**
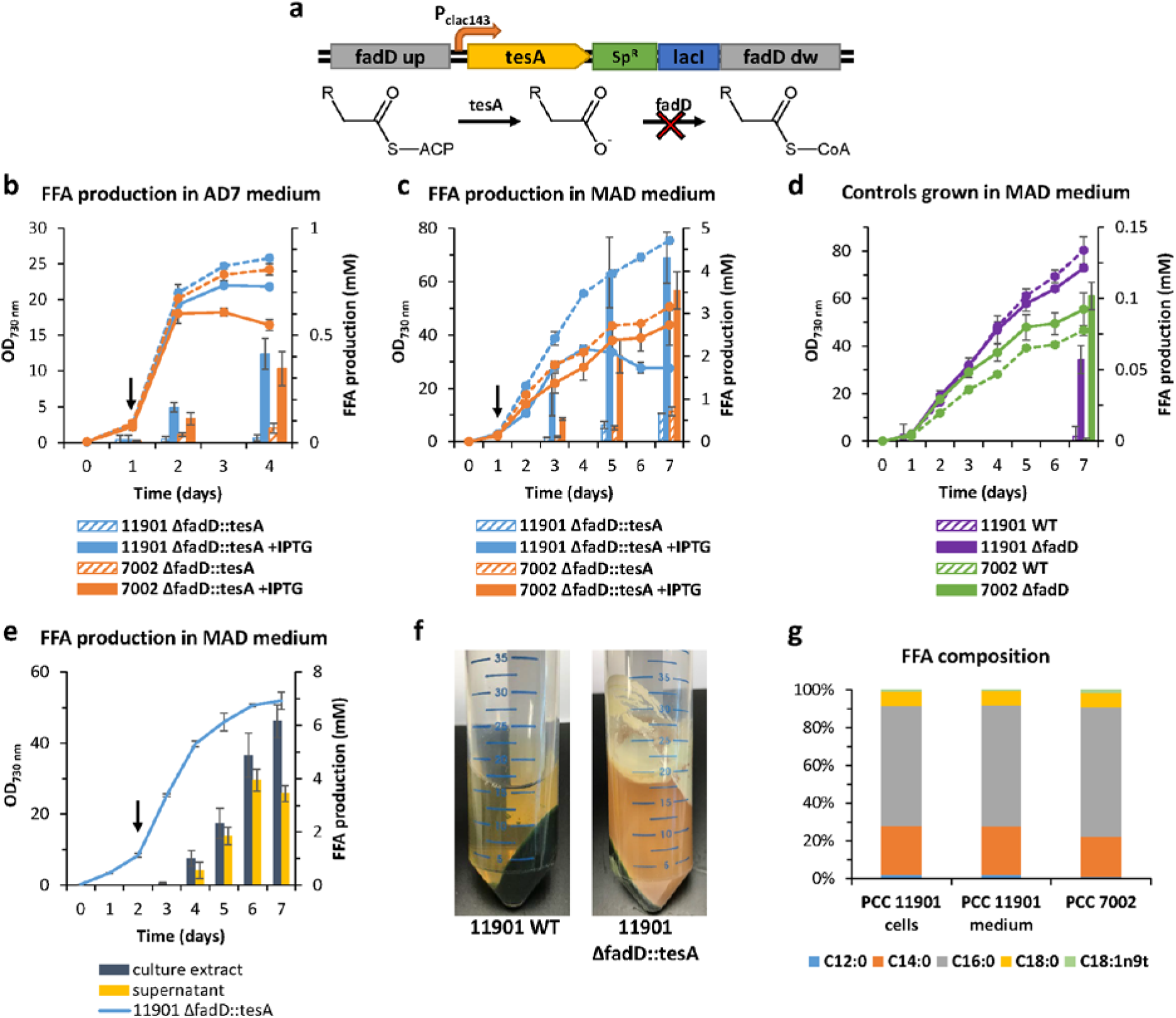
FFA production in engineered *Synechococcus* sp. PCC 11901 and 7002 strains. **(a)** Schematic illustration of FFA production pathway. The truncated acyl-CoA thioesterase gene (‘*tesA*) under control of P_clac143_ inducible promoter was inserted into chromosome with simultaneous knockout of the endogenous long-chain-fatty-acid--CoA ligase (*fadD*). ‘Tes A converts fatty acyl-ACP to free fatty acid and its reconversion to fatty acyl-CoA is negated by knocking out the endogenous *fadD*. Comparison of the growth and FFA productivity of the engineered *Synechococcus* sp. PCC 11901 and 7002 strains using regular AD7 **(b)** and MAD **(c)** medium. Filled and dashed lines indicate growth (OD_730_ measurements) of the induced and non-induced cultures respectively, while the column bars correspond to the FFA production. Arrows indicate addition of IPTG inducer and light intensity increase. **(d)** Comparison of growth and FFA production by the WT (dashed lines and columns) and Δ*fadD* knockout control (filled lines and columns) strains. **(e)** Analysis of FFA production by the engineered 11901 strain induced at OD_730_ ≈8. FFA were extracted either directly from the culture or from the medium supernatant. For OD_730_ measurements data points with error bars represent mean of n=3 biological replicates ± standard deviation (for 11901 ΔfadD::tesA strain n=2 in panels **b** and **c** and n=3 in panel **e**). FFA concentration as mean of n=6 replicates (3 biological and 2 technical) ± standard deviation (for 11901 ΔfadD::tesA strain n=4 in panels **b** and **c** and n=6 in panel **e**). **(f)** Detail of WT and engineered PCC 11901 strain cultures centrifuged after 7 days of cultivation. **(g)** Lipidomic profiles of the FFA from PCC 11901 and 7002 FFA producer strains: C12:0 (lauric acid), C14:0 (myristic acid), C16:0 (palmitic acid), C18:0 (stearic acid), C18:1n9t (elaidic acid).

Both 11901 and 7002 Δ*fadD*::*tesA* strains grown in AD7 medium produced similar amounts of FFA (0.41 and 0.34 mM respectively) 3 days after induction (Fig. 6b), while the non-induced strains produced less than 0.07 mM of FFA. The difference in productivity became much more apparent when the strains were grown in the MAD medium. The engineered 11901 Δ*fadD*::*tesA* strain was able to grow faster than the 7002 counterpart and 4 days after IPTG induction had already excreted 3.97 mM of FFA, a yield almost twice that achieved by the 7002 Δ*fadD*::*tesA* strain in the same time frame (2.01 mM) (Fig. 6c). While the 7002 Δ*fadD*::*tesA* strain was able to continue growing for two more days, achieving final FFAs yields of 3.54 mM, the corresponding 11901 strain declined and yields did not increase significantly after the 4^th^ day, reaching only 4.32 mM. When taking into account cell growth, the overall FFA productivity was higher for the 11901 strain (0.12 mmol/L/OD on day 5) compared to 7002 (0.05 mmol/L/OD and 0.08 mmol/L/OD on day 5 and 7 respectively). The control strain 7002 Δ*fadD* grown in optimized medium produced 0.1 mM of FFA, almost twice as much as the PCC 11901 control strain and production levels in the WT were negligible, within statistical error (Fig. 6d). The large error bars in all FFA measurements can be explained by the low solubility of FFA in water, which makes them float on top of the culture surface and adhere to the surface of culture vessels (Fig. 6f).

To test whether production yields could be further improved by using different extraction methods, the 11901 Δ*fadD*::*tesA* strain was grown in MAD medium and FFA were extracted with hexane either directly from cell cultures or from the cell-free medium supernatant (Fig. 6e). For the first few days, the difference between concentration of FFA in the cell culture and medium extracts was statistically insignificant. However, after 7 days, samples extracted directly from the cell culture contained significantly more FFA (6.16 mM) than the ones from the supernatant.

The composition of produced FFA was evaluated by Gas Chromatography analysis in both 11901 and 7002 Δ*fadD*::*tesA* engineered strains (**Fig. S9**). Despite having different final productivities, there was no major difference in the lipidomic profile of the two strains. For the 11901 strain, palmitic acid (C16:0) constitutes on average approximately 65% of the total FFA produced, followed by myristic acid (C14:0) – 23% and stearic acid (C18:0) – 9% (Fig. 6g).

## 3. Discussion

In this work we report the discovery of a novel cyanobacterial strain, *Synechococcus* sp. PCC 11901, from the Johor Strait (Singapore) and describe a new medium for its cultivation. PCC 11901 displays fast growth, high biomass accumulation and resistance to various abiotic stresses (such as high light and salinity), which are all important criteria for potential industrial uses of any cyanobacterial strain, especially when cultivated outdoors in a non-sterile environment and exposed to contamination that could potentially overgrow it. It was recently shown that engineering cyanobacteria to utilize unconventional phosphorous and nitrogen sources can dramatically reduce the risk of contamination^36^. However, increasing salinity of the growth medium is a more conventional method to decrease viability of less adaptable competing strains. We show that this euryhaline strain tolerates up to 10% (w/v) of NaCl and can grow at high light intensities greater than 750 µmol photons·m^-2^·s^-1^ (the limit of our equipment). Although PCC 11901 tolerates high temperatures, up to a maximum of 43 °C, the optimal temperature for its growth may be around 30 °C, the average temperature in its natural environment in Singapore with low fluctuations (28-32 °C) throughout the year. Though its cobalamin auxotrophy may be inconvenient for industrial cultivation, this can easily be overcome by heterologous expression of a cobalamin-independent methionine synthase (MetE)^24^. Interestingly, PCC 11901 strain already possesses a *metE* gene in one of the endogenous plasmids, though it seems to be inactive. This is possibly an evolutionarily recent mutation, most likely due to the ready availability of cobalamin in seawater^21^.

In our hands, PCC 11901 shows a similar doubling time to PCC 7002 of 2.14 ± 0.06 hours, slightly longer than that for UTEX 2973 (1.93 hours) (**Table S1**). However, compared to both UTEX 2973 and PCC 7002, PCC 11901 can grow to much higher cell densities and accumulate more biomass overall (Fig. 5a-b, **S4 and S6)**.

Experimental testing of nutrient limitations allowed us to tap into an unprecedented potential for high biomass accumulation. We show that PCC 11901 can grow to OD_730_ of 101 and produce 18.3 g/L of dry weight biomass, which are the highest reported values for any cyanobacterium. When grown alongside PCC 7002, PCC 7942, PCC 6803 and UTEX 2973 strains, PCC 11901 accumulated 2-3 more biomass than these strains despite using the same nutrient concentrations. It is also possible that PCC 11901 could produce even more biomass if exposed to greater light irradiances than we could provide, especially given that residual phosphate and iron precipitates were still present in the medium and no bleaching or pigment loss was observed even at the highest cell density. Growing marine strains in both MAD and the previously described MMA medium^31^ at higher temperatures (37-38 °C), led to a decline in biomass after 10 days of cultivation (**Fig. S6**). In contrast, when grown at 30 °C, strains could sustain growth to higher densities, possibly due to the higher solubility of CO_2_ at lower temperatures or a hitherto unknown regulatory mechanism. A comparison of the compositions of MAD and MMA media suggests that cyanobacteria do not require such a high excess of nutrients as found in MMA (especially phosphate, which is approximately 26x greater in MMA) to accumulate high biomass.

As a biotechnological chassis strain, PCC 11901 has several attractive traits: it is naturally transformable, which facilitates genetic manipulation; synthetic biology tools already existing for the PCC 7002 strain^32, 33^ are also compatible with this new strain, facilitating an easy switch of metabolic engineering constructs to the PCC 11901 strain and it shows high growth rates and high rates of biomass production. As a proof of concept, we expressed a truncated ‘*tesA* gene in order to produce FFA, a valuable biodiesel feedstock. FFA yields per litre of culture using MAD medium were found to be approximately 10-fold higher in comparison to using the regular AD7 medium under the same growth and induction conditions, even though the final OD_730_ was only 1.5 times higher. Our results demonstrate that growth medium optimization can dramatically increase the production yields of FFA though further investigations on the mechanism behind the observed changes to productivity will be necessary to further understand this phenomenon.

The maximum FFA titre achieved in this study of 6.16 mM (≈1.54 g/L, 5 days after induction) is the highest reported to date for cyanobacteria, with values of 0.64 g/L (when using isopropyl myristate overlay for FFA removal)^13^, 0.19 g/L^37^ and 0.13 g/L^12^ previously reported for PCC 7942, PCC 6803 and PCC 7002 respectively after 16-20 days of cultivation. It is also on a par with the production levels achieved by heterotrophic organisms. Introduction of different heterologous ‘*tesA* genes in a *ΔfadD* background strain of *E. coli* resulted in the production of ≈2 g/L FFA 2 days post-induction^38^ and impairment of glycogen metabolism in *Yarrowia lipolytica* led to production of 1.44 g/L of FFAs (up to 2.04 g/L when using a dodecane overlay) after 5 days^39^. Though these values are much less than the highest ever reported FFA production titre of 33.4 g/L, using heavily engineered *Saccharomyces cerevisiae* strains, this particular system required supplementation with >300 g/L of glucose over 10 days of cultivation^40^ and it is quite likely that further engineering of our strain will increase titre and cell viability.

Overall this study shows that the newly discovered cyanobacterium *Synechococcus* sp. PCC 11901, in conjunction with the use of modified growth medium, has the potential to become a relevant industrial biotechnology platform for the sustainable production of carbon-based molecules.

## 4. Materials and methods

### 4.1. Strain isolation and purification

PCC 11901 strain was isolated from the Johor Strait in the vicinity of Pulau Ubin island (Singapore, 1°25’17.7″N 103°57’20.6″E) and Johor river estuary (where the Johor river mixes with the seawater from the Singapore Strait and South China Sea). Water samples were collected at depth of 20 cm below the water surface into 50 mL falcon tubes, mixed with AD7 growth medium (1:1) for marine cyanobacteria (for details see Table S2) and transferred to culture flasks. Inoculums were grown at 38°C, 1% CO_2_, 160 rpm, under continuous illumination 200 µmol photons·m^-2^·s^-1^ for several days and subcultured three times. 50 µL of 2·10^5^ culture dilutions were plated on solid growth medium. To avoid exposing cells to potential mutagens (such as antibiotics commonly used to remove bacterial contamination), the enriched cultures were dilution-plated and resulting single colonies were consecutively re-streaked on solid AD7 medium. Single colonies were subsequently restreaked and xenic strains were additionally restreaked on plates containing cobalamin in order to remove bacterial contaminant. To check the purity of the new isolate, the strain was restreaked on LB agar plate and incubated for several days at 38°C. Both isolated strain and bacterial contaminant were identified as previously described by amplification of 16S rRNA fragments and Sanger sequencing using primer pairs CYA361f, CYA785r^41^ and 27f, 1492r^42^.

### 4.2. Strains and growth conditions

Newly isolated *Synechococcus* sp. PCC 11901 and PCC 7002 (obtained from the Pasteur Culture Collection) were grown photoautotrophically in medium AD7 (medium A^19^ with D7^20^ micronutrients lacking NaVO_3_). Solid medium was prepared by adding 1.2% (w/v) Bacto-Agar (BD Diagnostics) and 1 g/L sodium thiosulfate.

Cyanobacterial natural transformation was performed by adding 0.5 µg of respective plasmids to 1 mL of culture at OD_730_ ≈0.1 and incubating overnight under the same conditions. Cells were transferred onto solid media supplemented with either 50 µg/mL spectinomycin, 100 µg/mL kanamycin or 100 µM acrylic acid, as required, grown for 4 days and restreaked twice to ensure complete segregation.

All cloning steps were performed using supercompetent *E. coli* cells (Stellar, TaKaRa), grown in LB medium at 37 °C, supplemented with appropriate antibiotics (100 µg/mL carbenicillin or 50 µg/mL spectinomycin).

For starter cultures, freshwater cyanobacteria *Synechococcus elongatus* UTEX 2973 (a kind gift from Prof. Poul Erik Jensen), PCC 7942 (a gift from Prof. Adrian Fisher, Cambridge University) and *Synechocystis* sp. PCC 6803 (obtained from the Pasteur Culture Collection) were grown in medium BG-11^19^.

For all growth curve experiments using regular AD7 and BG-11 media, 25 mL of cultures adjusted to OD_730_ ≈0.1 were grown in Erlenmeyer baffled flasks, in triplicates, and shaken at 225 rpm using an NB-101SRC orbital shaker (N-Biotek, Korea) in a 740-FHC LED incubator (HiPoint Corporation, Taiwan), under constant illumination using an RGB LED Z4 panel set with R:G:B ratios either 4:2:1 (for cultures grown at 100, 300 and 500 µmol photons·m^-2^·s^-1^ light intensities) or 1:0:1 (for cultures grown at 660 µmol photons·m^-2^·s^-1^ light intensity). Lighting conditions were measured using a quantum flux meter (Apogee Instruments, model MQ-500), and temperature and CO_2_ conditions were set to values specified in **Table S1**, as required. Cells growth was monitored by measuring the optical density at 730 nm (OD_730_) in a 1-cm light path with a Cary 300Bio (Varian) spectrophotometer. Doubling times were calculated as mean within the logarithmic range only.

For the cobalamin auxotrophy experiment, a starter culture of the axenic strain was grown in medium AD7 supplemented with cobalamin, but the inoculum was washed 3 times in order to remove any remaining cobalamin from the medium.

To facilitate acclimation to high salt concentrations cultures were incubated in low light for 1 week. Once the cultures were adapted to the respective conditions, biological triplicates were inoculated to starting OD_730_ of 0.1 and grown at 38 °C and 300 µmol photons·m^-2^·s^-1^ light intensity.

For the biomass evaluation in the AD7 medium optimization experiment 0.4 to 1 mL of cell cultures were filtered onto pre-dried and pre-weighed glass microfiber filters (47 mm diameter, 1 µm pore size, GE Healthcare, Cat. No. 1822-047), washed three times with MilliQ water in order to remove salt remains and dried for 24 hours at 65 °C (at which point measured weight did not differ by more than ± 0.0001 g).

For the comparison of all cyanobacterial strains 3x 33 mL of either MAD or 5xBG (for details see **Table S2**) media were inoculated with cells to starting OD_730_ of 0.1 and grown with shaking at 180 rpm, 30 °C, with RGB LED ratio 1:1:1. For growth of PCC 11901, PCC 7002 and UTEX 2973, the initial light intensity was set to 150 µmol photons·m^-2^·s^-1^ increased after 1 day to 750 µmol photons·m^-2^·s^-1^. In the case of PCC 7942 and PCC 6803 the initial light intensity was set to 75 µmol photons·m^-2^·s^-1^, changed to 150 µmol photons·m^-2^·s^-1^ after 1 day and further increased to 750 µmol photons·m^-2^·s^-1^ on the next day. To compensate for water evaporation 700 µL of sterile MilliQ water was added to the cultures on daily basis. For dry biomass evaluation, 0.5 to 1 mL of cultures were transferred into pre-weight Eppendorf tubes and centrifuged at 20,000 × g for 2 minutes. Cell pellets were washed three times with 1 mL of MilliQ water in order to dissolve and remove remaining salt precipitates and dried at 65 °C for 24 hours.

### 4.3. Genome sequencing

Genomic DNA was extracted using Quick-DNA Fungal/Bacterial Kit (Zymo Research), concentrated and analysed by PacBio and Illumina MiSeq next-generation sequencing. Illumina MiSeq sequencing library construction was performed according to Illumina’s TruSeq Nano DNA Sample Preparation protocol.

Libraries were pooled at equimolar concentrations and sequenced on the Illumina MiSeq platform at a read-length of 300 bp paired-end.

PacBio run library preparation was performed using SMRTbell template prep kit 1.0 (Pacific Biosciences, USA) followed by single-molecule real-time sequencing on the PacBio RS II platform. The PacBio reads were assembled using HGAP3 assembler^43^, and then polished with Quiver within the SMRT Analysis v2.3.0 protocol. Polished assembly contigs were then circularized and re-oriented with Circlator 1.1.4^44^. Assembled genomes were corrected by mapping 3 MiSeq Illumina paired reads to the reference sequences using the Geneious Prime^®^ 2019.1.3 software (Biomatters Ltd.) and consensus sequences were exported and deposited to GenBank (accession number CP040360.1). Average nucleotide identity (ANI) analysis was performed using OrthoANIu algorithm^45^. Circular diagrams of the PCC 11901 genome sequence and BLAST analysis were created using CGView Server^BETA^ online tool^28^.

### 4.4. Glycerol and glucose adaptation

AD7 medium supplemented with 10 mM glycerol or 0.15% (w/v) glucose was inoculated with a single colony of PCC 11901 and grown for one week at low light intensity 50 µmol photons·m^-2^·s^-1^, 38°C, 1% CO_2_. Growth was observed after a few days and strains were restreaked on AD7 agar plates with either 10 mM glycerol or 0.15% (w/v) glucose. For the photoheterotrophy assay glycerol and glucose adapted strains were grown to OD_730_ ≈5 and 15 µL dilution series were transferred onto AD7 agar plates containing different combinations of 10 µM DCMU, 10 mM glycerol or 0.15% (w/v) glucose. Plates were dried and incubated for 7 days at 50 µmol photons·m^-2^·s^-1^, 38°C, 1% CO_2_.

### 4.5. Cloning of constructs

All fragments needed for cloning of pSW036, pSW039, pSW040, pSW068 and pSZT025 vectors were amplified using Q5^®^ High-Fidelity DNA Polymerase (New England Biolabs) according to manufacturer’s protocol. All primers and templates used for PCR amplification are listed in **Table S2**. PCR products were incubated overnight with Dpn*I* restriction enzyme and purified using an EZ-10 Spin Column PCR Products Purification Kit (BioBasic). Fragments were then ligated using NEBuilder^®^ HiFi DNA Assembly (New England Biolabs) according to manufacturer’s protocol and transformed to competent *E. coli* cells (Stellar, TaKaRa). A synthetic version of the *E. coli* thioesterase gene *tesA* lacking its cognate signal sequence (*‘tesA*) was codon-optimized for Syn7002 (by GenScript, Hong Kong, Ltd). pAcsA_cpt_YFP and pAcsA_cLac143_YFP^32^ vector templates for PCR were a kind gift from Prof. Brian Pfleger, University of Wisconsin-Madison, USA.

### 4.6. YFP fluorescence measurements

Triplicates of WT, Δ*acsA*::P_cpt_-YFP and Δ*psbA2*::P_clac143_-YFP strains were grown in regular liquid AD7 medium to OD_730_ ≈1 (time 0), and, if required, induced with 1 mM IPTG. Whole cell fluorescence was measured by transferring 150 µL of cultures, collected at T=0, after 6 and 24 hours into 96-well black clear bottom plates and measured with a Hidex Sense plate reader, using 485/10 and 535/20 nm filters for excitation and emission, respectively. Relative fluorescence was normalized to OD_730_ (RFU/ OD_730_). All measurements were performed in triplicates.

### 4.7. Fluorescence microscopy

Liquid cultures of WT, Δ*acsA*::P_cpt_-YFP and Δ*psbA2*::P_clac143_-YFP strains were harvested at OD_730_ ≈1 by centrifugation, concentrated and transferred onto 0.5% agarose pads placed on microscopy slides. Once dry, pads were covered with a coverslip. Fluorescence microscopy was performed using Axio Observer Z1 (Zeiss) inverted fluorescence microscope with EC Plan-Neofluar 100x/1.30 Oil Ph 3 objective and immersion oil (total magnification 1000x). For the chlorophyll autofluorescence and YFP fluorescence excitation/emission wavelengths were 577/603 and 525/538 nm respectively and images were exposed for 130 ms.

### 4.8. Transmission electron microscopy

Cultures were harvested by centrifugation, fixed for 1 h at room temperature with 4% (w/v) glutaraldehyde in 100 mM phosphate buffer (pH 7.3) and washed 3× with 100 mM phosphate buffer. After embedding in 2% (w/v) low-gelling-temperature agarose, samples were cut in 1–2 mm cubic blocks, and post-fixed with 1% (w/v) osmium tetroxide in distilled water for 1 h. Samples were washed twice with distilled water, and dehydrated through a graded ethanol series (1 × 15 min 50%, 1 × 15 min 70%, 1 × 15 min 90% and 3 × 20 min 100%). Two 5 min washes with acetone were performed prior to infiltration with araldite for 1 h and with fresh Araldite overnight. Polymerisation was achieved by incubation at 60–65°C for 48 h. Ultrathin sections were cut with a diamond knife at a Reichert Ultracut E microtome and collected on uncoated 300-hexagonal mesh copper grids (Agar Scientific). High contrast was obtained by post-staining with saturated aqueous uranyl acetate and Reynolds lead citrate for 4 min each.

Negative staining was performed on 300-mesh copper carbon supports grids (Agar Scientific) that were previously rendered hydrophilic by glow discharge (Easy-Glow, Ted Pella). Glutaraldehyde fixed bacteria were adsorbed to TEM grids by direct application of 5 µl of the suspension for 1 minute and stained by floating the loaded grid onto a drop of 1% uranyl acetate for 20 seconds. The grids were examined in a JEOL JEM-1230 transmission electron microscope at an accelerating potential of 80 kV.

### 4.9. Free fatty acid production assay

PCC 11901 strain was transformed with the pSW068 and pSW071 vectors generating 11901 Δ*fadD*::*tesA* (production) and Δ*fadD* (control) strains respectively. PCC 7002 was transformed with pSZT025 and pSW072 to generate 7002 Δ*fadD*::*tesA* (production) and Δ*fadD* (control) strains. Successful transformants were screened for complete segregation by colony PCR (**Fig. S10**). A large number of the screened colonies for the PCC 11901 strain did not carry the designed insert, suggesting that kanamycin may not be the antibiotic of choice for selection in this strain, as it exhibits partial resistance to kanamycin at low concentrations.

All engineered and control strains were grown in 33 mL cultures, in biological triplicates, (except for PCC 11901 Δ*fadD*::*tesA* grown in biological duplicates) using either basic AD7 or MAD medium. Cultures were inoculated with cells to starting OD_730_ of 0.1 and grown with shaking at 180 rpm, 30 °C, with RGB LED ratio 1:1:1 at 150 µmol photons·m^-2^·s^-1^ light intensity. After 1 day, cultures were induced with 1 mM IPTG and the light was increased to 750 µmol photons·m^-2^·s^-1^. 1 mL medium aliquots were collected for both OD_730_ and FFA quantification. For FFA quantification, cell cultures were centrifuged for 2 minutes at 20,000 × g and medium supernatants were carefully collected to avoid any disruption of the cell pellets. FFA were quantified in technical duplicates using the EZScreen™ Free Fatty Acid Colorimetric Assay Kit (384-well) (BioVision, USA) according to manufacturers’ protocol.

For the GC analysis both cell extract and medium supernatant were acidified with 1 M HCl to pH ≈ 2 in order to protonate FFA and facilitate extraction. Samples were extracted with *n*-hexane, evaporated and dried using a centrifuge vacuum concentrator. Dried samples were resuspended in 100 µL of hexane and aliquots were transferred on the TLC Silica gel 60 plates (Merck, Germany). Plates were resolved for approximately 30 minutes in hexane, diethyl ether, formate solvent mixture at the 70:30:2 ratio. Sample preparation and the GC analysis was performed as previously described^46^.

## Abbreviations

FFA: free fatty acids
OD_730_: optical density measured at 730 nm wavelength
gDW: grams of dry weight
DCMU: 3-(3,4-dichlorophenyl)-1,1-dimethylurea
RGB LED: red green blue light-emitting diode

## Contributions

AW and TTS performed all experiments. Conception, design and data analysis were carried out by AW, TTS and PJN. Manuscript was written by AW, TTS and PJN. BN critically reviewed the manuscript.

## Competing interests

The authors declare no competing interests.

## Acknowledgements

This work was supported by NTU grants M4080306 to BN and M4081714 to PJN. The authors would like to thank Daniela Moses (SCELSE, NTU) for assistance with whole genome sequencing reactions, Anthony Wong (SCELSE, NTU) for providing access to the fluorescence microscopy facility, Giulia Mastroianni (School of Biological and Chemical Sciences, Queen Mary University of London) for TEM sample preparation and analysis and Michael Voigtmann (Wintershine Pte. Ltd.) for providing access to the sample collection site. The authors are also grateful to Prof. Bertil Andersson for his constant support and encouragement and his pivotal role in establishing the CyanoSynBio@NTU laboratory.

