## Supplementary material for "Unprecedented biomass and fatty acid production by the newly discovered cyanobacterium *Synechococcus* sp. PCC 11901"

**Table S1.** Doubling times of *Synechococcus* sp. PCC 11901, *Synechococcus* sp. PCC 7002 and *Synechococcus elongatus* UTEX 2973. Strains were grown side by side under different conditions. For all lighting conditions RGB ratio was set to 4:2:1 except for 660*, where the red and blue light were used only at 1:1 ratio. All doubling times are given in hours. Average doubling time was calculated as mean of n=3 biological replicates ± standard deviation.

| | $0.04\%\ CO_2$ | | $1\%\ CO_2$ | | | | | |
|---|---|---|---|---|---|---|---|---|
| **Temp. (°C)** | **38** | | **38** | | | **41** | | **30** |
| **Light intensity (µmol photons·m$^{-2}$·s$^{-1}$)** | **100** | **300** | **300** | **500** | **660*** | **300** | **500** | **300** |
| **PCC 11901** | 3.35 ± 0.12 | 3.85 ± 0.12 | 2.46 ± 0.02 | 2.33 ± 0.02 | 2.14 ± 0.06 | 2.62 ± 0.07 | 2.80 ± 0.08 | 3.20 ± 0.05 |
| **PCC 7002** | 3.59 ± 0.12 | 3.96 ± 0.22 | 2.46 ± 0.09 | 2.27 ± 0.01 | 2.29 ± 0.13 | 2.55 ± 0.13 | 2.83 ± 0.16 | 2.97 ± 0.06 |
| **UTEX 2973** | 3.65 ± 0.32 | 3.05 ± 0.05 | 2.08 ± 0.06 | 2.02 ± 0.01 | 2.02 ± 0.04 | 2.15 ± 0.05 | 1.93 ± 0.04 | 3.09 ± 0.10 |

**Table S2.** Media formulations used for the growth experiments. **All concentrations are in mM**. Media in which marine (MAD) and freshwater (5xBG) cyanobacterial strains performed best were marked with star (*).

| | AD7 | BG-11 | MAD* | MBG | MBG-Mg+ | 5xBG* | 5xBGM |
|---|---|---|---|---|---|---|---|
| $NaNO_3$ | 12 | 17.6 | 96 | 96 | 96 | 88 | 96 |
| $KH_2PO_4$ | 0.37 | - | 1.2 | - | - | - | 0.6 |
| $K_2HPO_4$ | - | 0.175 | - | 1.2 | 1.2 | 0.875 | 0.6 |
| NaCl | 308 | - | 308 | - | - | - | - |
| KCl | 8 | - | 8 | - | - | - | - |
| $CaCl_2 \cdot 2H_2O$ | 2.5 | 0.245 | 2.5 | 0.245 | 0.245 | 1.22 | 1.22 |
| $Na_2EDTA$ | 0.081 | 0.027 | 0.081 | 0.027 | 0.027 | 0.13 | 0.13 |
| $MgSO_4 \cdot 7H_2O$ | 20.3 | 0.3 | 20.3 | 0.3 | 20.3 | 1.52 | 1.52 |
| $FeCl_3 \cdot 6H_2O$ | 0.014 | - | 0.24 | - | - | - | - |
| Ammonium iron (III) citrate | - | 0.023 | - | 0.24 | 0.24 | 0.24 | 0.24 |
| Tris-HCl | 8.6 | - | 8.6 | - | - | - | - |
| $H_3BO_3$ | 0.046 | 0.046 | 0.046 | 0.046 | 0.046 | 0.231 | 0.231 |
| $MnCl_2 \cdot 4H_2O$ | 0.0091 | 0.0091 | 0.0091 | 0.0091 | 0.0091 | 0.0457 | 0.0457 |
| $ZnSO_4 \cdot 7H_2O$ | 0.00077 | 0.00077 | 0.00077 | 0.00077 | 0.00077 | 0.00386 | 0.00386 |
| $Na_2MoO_4 \cdot 2H_2O$ | 0.00521 | 0.00161 | 0.00521 | 0.00161 | 0.00161 | 0.00806 | 0.00806 |
| $CuSO_4 \cdot 5H_2O$ | 0.00032 | 0.00032 | 0.00032 | 0.00032 | 0.00032 | 0.00158 | 0.00158 |
| $(CoNO_3)_2 \cdot 6H_2O$ | - | 0.00017 | - | 0.00017 | 0.00017 | 0.00084 | 0.00084 |
| $CoCl_2 \cdot 6H_2O$ | 0.00017 | - | 0.00017 | - | - | - | - |
| vitamin $B_{12}$ | 0.000003 | - | 0.000003 | - | - | - | - |
| $Na_2CO_3$ | - | 0.189 | - | 0.189 | 0.189 | 0.943 | 0.943 |
| citric acid | - | 0.031 | - | 0.031 | 0.031 | 0.156 | 0.156 |

**Table S3.** List of vectors and primers used for the amplification of DNA blocks for the Gibson assembly.

| Plasmid name | PCR template | PCR product | Primer name | Primer sequence (5’-3’) |
|---|---|---|---|---|
| pSW036 | Synechococcus sp. PCC 11901 gDNA | acsA upstream FR | SSW07_acsA_up_F | GACGTTGTAAAACGACGGCCAGTgcaatgctgagatgatcctcggtgaa |
| | | | SSW07_acsA_up_R | ttattatcgtgggatttatttcaccccatt |
| | pAcsA_cpt_YFP* | Pcpt-YFP | cpt-YFP_acsA_F | aatggggtgaaataaatcccacgataataattaacaaaaaagcaggaataaaattaacaagatgtaac |
| | | | cpt-YFP_acsA_R | aagacaccctctgtcctctggacatctttgaggccgttgatctagacaaa |
| | Synechococcus sp. PCC 11901 gDNA | acsA downstream FR | SSW07_acsA_dw_F | agatgtccagaggacagagggtgt |
| | | | SSW07_acsA_dw_R | CAATTTCACACAGGAAACAGCTATGACagtaaatcagagacagaaaccttccagacg |
| | pUC19 | pUC backbone | pUC19_B_F | GTCATAGCTGTTTCCTGTGTGAAATTGTTATC |
| | | | pUC19_B_R | ACTGGCCGTCGTTTTACAACGT |
| pSW039 | Synechococcus sp. PCC 11901 gDNA | psbA2 upstream FR | SSW07_psbA2_up_F | CAGTCACGACGTTGTAAAACGACGGCCAGTaattagtgtaggagatcaccgttgaattgatcac |
| | | | SSW07_psbA2_up_R | aattgcatgtaccagtttaaagtactgggg |
| | pAcsA_cLac143_YFP* | Pclac143-YFP | clac143-YFP_psbA2_F | ccccagtactttaaactggtacatgcaattgactcccctctggacatctcca |
| | | | clac143-YFP_psbA2_R | aaccacacgcacaattcccttaaaaagcaacattaattgcgttgcgctcactg |
| | Synechococcus sp. PCC 11901 gDNA | psbA2 downstream FR | SpR_psbA2_F | tgctttttaagggaattgtgcgtgtggtt |
| | | | SpR_psbA2_R | ccactgcaatcccaaattcaaacttccagAAACGGATGAAGGCACGAACCCAG |
| | pDF-trc* | $\text{Spec}^R$ | SSW07_psbA2_dw_F | ctggaagtttgaatttgggattgcagtgg |
| | | | SSW07_psbA2_dw_R | TTTCACACAGGAAACAGCTATGACtggcaggcgagacaactgctttcg |
| | pUC19 | pUC backbone | pUC19_B_F | GTCATAGCTGTTTCCTGTGTGAAATTGTTATC |
| | | | pUC19_B_R | ACTGGCCGTCGTTTTACAACGT |
| pSZ013 (used for subcloning only) | codon optimized tesA from GenScript | tesA | ICA_tesA_to_cLac_F | gataacaatttcacacaccaactcataaagtcaagtaggagattaattccATGGATACCCTCCTCATTCTC |
| | | | ICA_tesA_to_cLac_R | gtggcagcagccaactcagcttcctttcgggctttgttagacagccggatAGGGCCGAGTTTGTACAAG |
| | pAcsA_cLac143_YFP | pAcsA_cLac143_YFP backbone | pAcsA_bef_pMB2_F | ATCCGGCTGTCTAACAAAG |
| | | | pAcsA_lacO_R | GGAATTAATCTCCTACTTGACTTTATG |
| pUC19-fadD (used for subcloning only) | codon optimized tesA from GenScript | tesA | InF_fadD_F | ACCCGGGGATCCTCTTCAGGTCAATGACATTGC |
| | | | InF_fadD_R | CTGCAGGTCGACTCTACCAGATTATCGCCCACTTTC |
| | pUC19 | puc19 backbone | XbaI digested | |
| pSZ025 | pUC57-kan | $\text{Kan}^R$ | ICA_KanR_F | gggatctcgaccgatgcccttgagaATTAAGGAGTGGACAacattgc |
| | | | ICA_KanR_R | TCCGCCATTCGGATCGCTTAATCTACGTGAACCAAGTCATTAGAAAAAC |

| | | | | |
|---|---|---|---|---|
| | pUC19 | puc19 backbone | ICA_open_fadD_F | gttttctaaTGACTTGGTTCACGTAGATTAAGCGATCCGAATGG |
| | | | ICA_open_fadD_R | CACAATTCGtttggagatgtccagaAATTCCATTTAAAAGCGCAAAAAAC |
| pSW040 (used for subcloning only) | Synechococcus sp. PCC 11901 gDNA | fadD FR | SSW fadD up F2 | GTAACGCCAGGGTTTTCCCAGTCACGACGTTGTAAAACGACGGCCAGTtaattaaaatttgactgcgatcgcctgcaatggcctgggcctgggcgacaat |
| | | | SSW07 fadD dw R2 | ATGTTGTGTGGAATTGTGAGCGGATAACAATTTCACACAGGAAACAGCTATGACaaatcgccgtggatttgaggtggggaataagggtttgagccgtg |
| | pUC19 | pUC backbone | pUC19_B_F | GTCATAGCTGTTTCCTGTGTGAAATTGTTATC |
| | | | pUC19_B_R | ACTGGCCGTCGTTTTACAACGT |
| pSW068 | pSZ025 | clac143-tesA | clac143-fadD F | cttttaaatggaattgcctctggacatctccaaaCGAATTGTGA |
| | | | clac143-fadD R | gatatatttttatcttgtgcaatgtTAATCTagAAAGATGACTAATTCGCTCGTC |
| | pSW040 | pUC-fadD backbone | pUC19_B_F | GTCATAGCTGTTTCCTGTGTGAAATTGTTATC |
| | | | pUC19_B_R | ACTGGCCGTCGTTTTACAACGT |
| pSW071 | pSW068 | pSW068 w/o tesA | SSW fadD KO F | tggaattgccacattgcacaagataaaaatatatcatcatgaacaataaaact |
| | | | SSW fadD KO R | caatgtggcaattccatttaaaagcgcaaaaaacaag |
| pSW072 | pSZ025 | pSZ025 w/o tesA | SSW fadD KO F | tggaattgccacattgcacaagataaaaatatatcatcatgaacaataaaact |
| | | | SSW fadD KO R | caatgtggcaattccatttaaaagcgcaaaaaacaag |

**Table S4**. List of primers used for sequencing of constructs and genotyping of transformed cyanobacterial strains.

| Primer name | Primer sequence (5’-3’) |
|---|---|
| CYA361f | GGAATTTTCCGCAATGGG |
| CYA785r | GACTACWGGGGTATCTAATCC |
| 27f | AGAGTTTGATCCTGGCTCAG |
| 1492r | TACCTTGTTACGACTT |
| SSW_seg_acsA_F | aggtcatatccgaggcgtacattca |
| SSW_seg_acsA_R | gaactaggtcaaggcaagcagtcgg |
| SSW07 psbA2 seg F | gtgattaagcaataaatcgattgagcga |
| SSW07 psbA2 seg R | tcaggattcagagcaaaccaagatta |
| fadD SSW07 seg F | agtaggattgtagccatgatttcgg |
| fadD SSW07 seg R | taggcacttggtttccgtccat |

**Table S5.** List of predicted proteins present in genome major insertions of *Synechococcus* sp. PCC 11901 found by BLAST search analysis.

| Locus tag | Protein name (highest identity) |
|---|---|
| FEK30_11785 | ABC transporter permease [Synechococcus sp. NKBG042902] |
| FEK30_11790 | ABC transporter ATP-binding protein [Synechococcus sp. NKBG042902] |
| FEK30_11795 | acyltransferase [Synechococcus sp. NKBG042902] |
| FEK30_11800 | FkbM family methyltransferase [Synechococcus sp. NKBG042902] |
| FEK30_11805 | glycosyltransferase family 4 protein [Synechococcus sp. NKBG042902] |
| FEK30_11810 | glycosyltransferase [Synechococcus sp. NKBG042902] |
| FEK30_11815 | glycosyltransferase family 2 protein [Nodularia sp. NIES-3585] |
| FEK30_11820 | glycosyltransferase [Nostoc sp. NIES-3756] |
| FEK30_11825 | hypothetical protein [Nodularia sp. NIES-3585] |
| FEK30_11830 | glycosyltransferase [Nodularia sp. NIES-3585] |
| FEK30_11835 | glycosyltransferase family 2 protein [filamentous cyanobacterium CCP2] |
| FEK30_11840 | glycosyltransferase [Nostoc sp. PCC 7524] |
| FEK30_11845 | glycosyltransferase family 2 protein [Synechococcus sp. NKBG042902] |
| FEK30_11850 | acyltransferase [Synechococcus sp. NKBG042902] |
| FEK30_11855 | glycosyltransferase family 2 protein [Nostoc sp. NIES-3756] |
| FEK30_11860 | glycosyltransferase family 4 protein [Synechococcus sp. NKBG042902] |
| FEK30_11865 | glycosyltransferase family 2 protein [Synechococcus sp. NKBG042902] |
| FEK30_11870 | glycosyltransferase family 4 protein [Synechococcus sp. NKBG042902] |
| FEK30_11875 | glycosyltransferase family 4 protein [Synechococcus sp. NKBG042902] |
| FEK30_11880 | hypothetical protein [Synechococcus sp. NKBG042902] |
| FEK30_11885 | glycosyltransferase [Synechococcus sp. NKBG042902] |
| FEK30_12515 | IS5 family transposase [Synechococcus sp. NKBG042902] |
| FEK30_12765 | DGQHR domain-containing protein [Nostocales cyanobacterium] |
| FEK30_13300 | DUF3854 domain-containing protein [Synechococcus sp. NKBG042902] |
| FEK30_13890 | phospholipid carrier-dependent glycosyltransferase [Synechococcus sp. PCC 7003] |
| FEK30_15055 | hypothetical protein [Trichormus sp. NMC-1] |
| FEK30_15475 | IS256 family transposase [Synechococcus sp. NKBG042902] |
| FEK30_15980 | F0F1 ATP synthase subunit beta [Synechococcus sp. NKBG042902] |
| FEK30_15515 | IS5 family transposase [Synechococcus sp. NKBG042902] |
| FEK30_15850 | IS5 family transposase [Synechococcus sp. NKBG042902] |
| FEK30_01650 | NAD(P)-dependent alcohol dehydrogenase [Synechococcus sp. NKBG042902] |
| FEK30_01865 | DNA cytosine methyltransferase [Coleofasciculus chthonoplastes] |
| FEK30_01875 | helix-turn-helix transcriptional regulator [Chroococcidiopsis sp. CCALA 051] |
| FEK30_03655 | BrnT family toxin [Anabaenopsis circularis] |
| FEK30_06150 | TPA: methicillin resistance protein [Syntrophomonas sp.] |
| FEK30_06155 | MULTISPECIES: methionyl-tRNA formyltransferase [Vibrio] |
| FEK30_06165 | translocase [Anaerolineaceae bacterium 4572_78] |
| FEK30_09280 | type II toxin-antitoxin system RelB/DinJ family antitoxin [Synechococcus sp. BDU 130192] |

| | |
|---|---|
| FEK30_09285 | type II toxin-antitoxin system YafQ family toxin |
| FEK30_09390 | MULTISPECIES: Fic family protein [Synechococcus] |
| FEK30_09715 | IS5 family transposase [Synechococcus sp. NKBG042902] |
| FEK30_09720 | MULTISPECIES: IS982 family transposase [unclassified Cyanobacteria (miscellaneous)] |
| FEK30_10375 | IS630 family transposase [Cyanothece sp. PCC 7425] |
| FEK30_10580 | MULTISPECIES: IS982 family transposase [unclassified Cyanobacteria (miscellaneous)] |
| FEK30_11580 | MULTISPECIES: S-layer homology domain-containing protein [Synechococcus] |

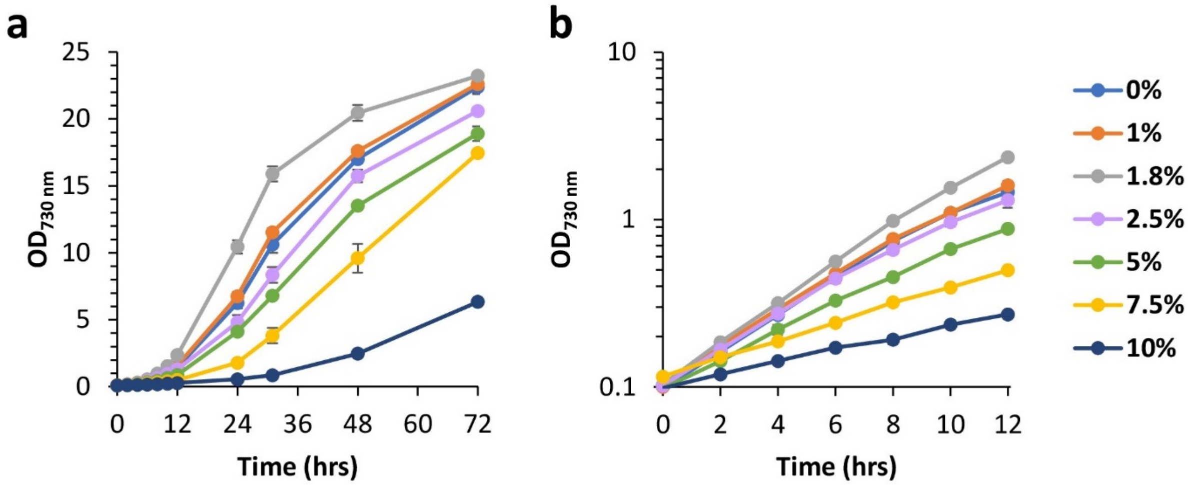


**Figure S1.** Salt tolerance analysis of PCC 11901 strain. **(a)** Cultures were grown in triplicates for 72 hours at 38 °C, 225 rpm shaking, constant illumination of 300 µmol photons·$m^{-2}$·$s^{-1}$ using AD7 media with varying sodium chloride (w/v) concentrations. **(b)** In the exponential growth phase, measurements were taken in 2-hours intervals. Time points within exponential phase were used for calculating the average growth rate. Average $OD_{730}$ was calculated as mean of n=3 biological replicates ± standard deviation.

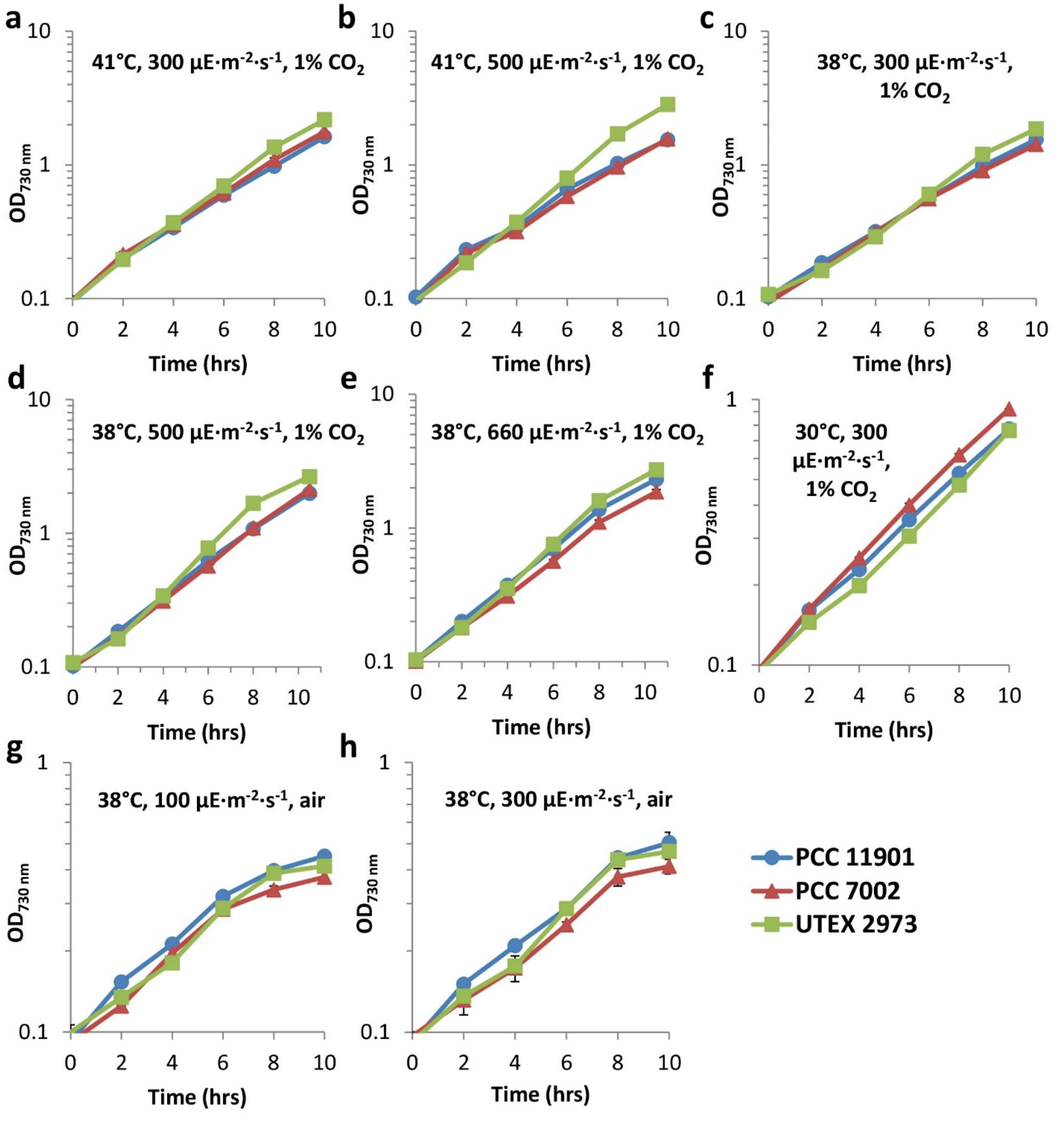


**Figure S2.** Growth performance comparison of PCC 11901, PCC 7002 and UTEX 2973 strains in different conditions. Strains, depending on requirements were grown in either AD7 or BG-11 medium, under constant illumination using LED RGB light 4:2:1 ratio setting **(a-d, f-h)**, except for growth curve **(e)** where red and blue LEDs only were used at 1:1 ratio. All strains were inoculated to initial $OD_{730}$ of approximately 0.1. Cultures were grown in triplicates and $OD_{730}$ measurements were taken in 2-hours intervals. Time-points within exponential growth phase only were used for calculating average doubling time. In blue growth curves of PCC 11901, in red PCC 7002 and in green UTEX 2973. Average $OD_{730}$ was calculated as mean of n=3 biological replicates ± standard deviation.

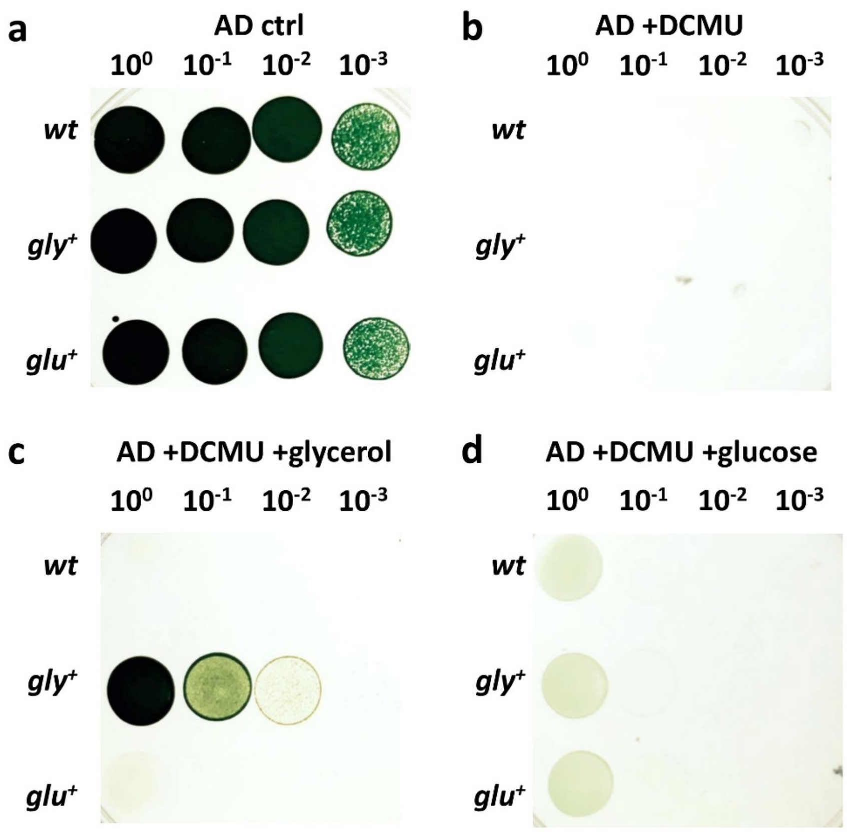


**Figure S3.** Glycerol and glucose tolerance of PCC 11901 and photoheterotrophy analysis. *wt*, glycerol and glucose adapted strains were grown to $OD_{730}$ = 5 and subsequently diluted. Dilutions were transferred onto solid AD7 medium **(a)** without any additives, **(b)** with 10 µM DCMU of herbicide DCMU, **(c)** 10 µM DCMU and 10 mM glycerol and finally **(d)** 10 µM DCMU and 0.15 % glucose.

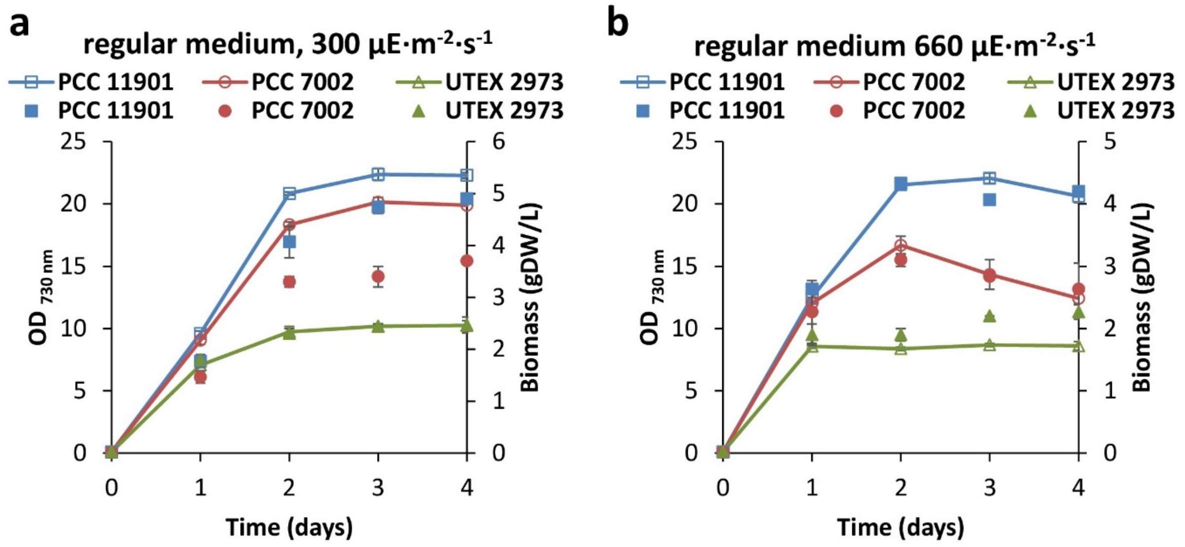


**Figure S4.** Growth performance and biomass accumulation of PCC 11901, PCC 7002 and UTEX 2973 using basic AD7 or BG-11 media. Lines with empty markers correspond to $OD_{730}$ measurements, whereas filled markers correspond to biomass accumulation. Strains were grown at 38 °C, 225 rpm at either **(a)** 300 µmol photons·m$^{-2}$·s$^{-1}$ RGB 4:2:1 or **(b)** 660 µmol photons·m$^{-2}$·s$^{-1}$ RB 1:1. Average $OD_{730}$ and biomass were calculated as mean of n=3 biological replicates ± standard deviation.

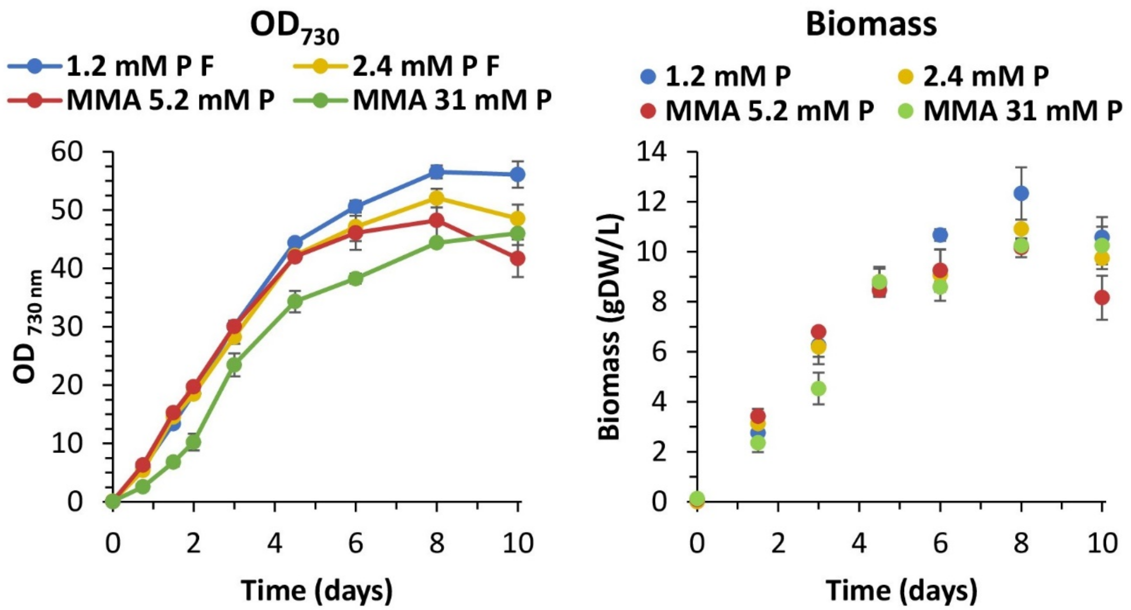


**Figure S5**. Comparison of PCC 11901 strain's growth in different modified media. Two enriched medium formulation variants used in this study were compared with previously published MMA medium optimized for 7002. All cultures were grown in triplicates at 38 °C, 225 rpm, 300 µmol photons·$m^{-2}$·$s^{-1}$. $OD_{730}$ and dry cell weight were measured in time intervals for 10 days. In blue AD medium supplemented with 96 mM $NaNO_3$, 1.2 mM $KH_2PO_4$ and 240 µM $FeCl_3$ (MAD), in yellow is the same medium but with more phosphate added (2.4 mM $KH_2PO_4$). Red and green markers correspond to MMA medium (120 mM $NaNO_3$, 1.1 mM $FeCl_3$) with either 5.2 or 31 mM $KH_2PO_4$ respectively. Average $OD_{730}$ and biomass were calculated as mean of n=3 biological replicates ± standard deviation.

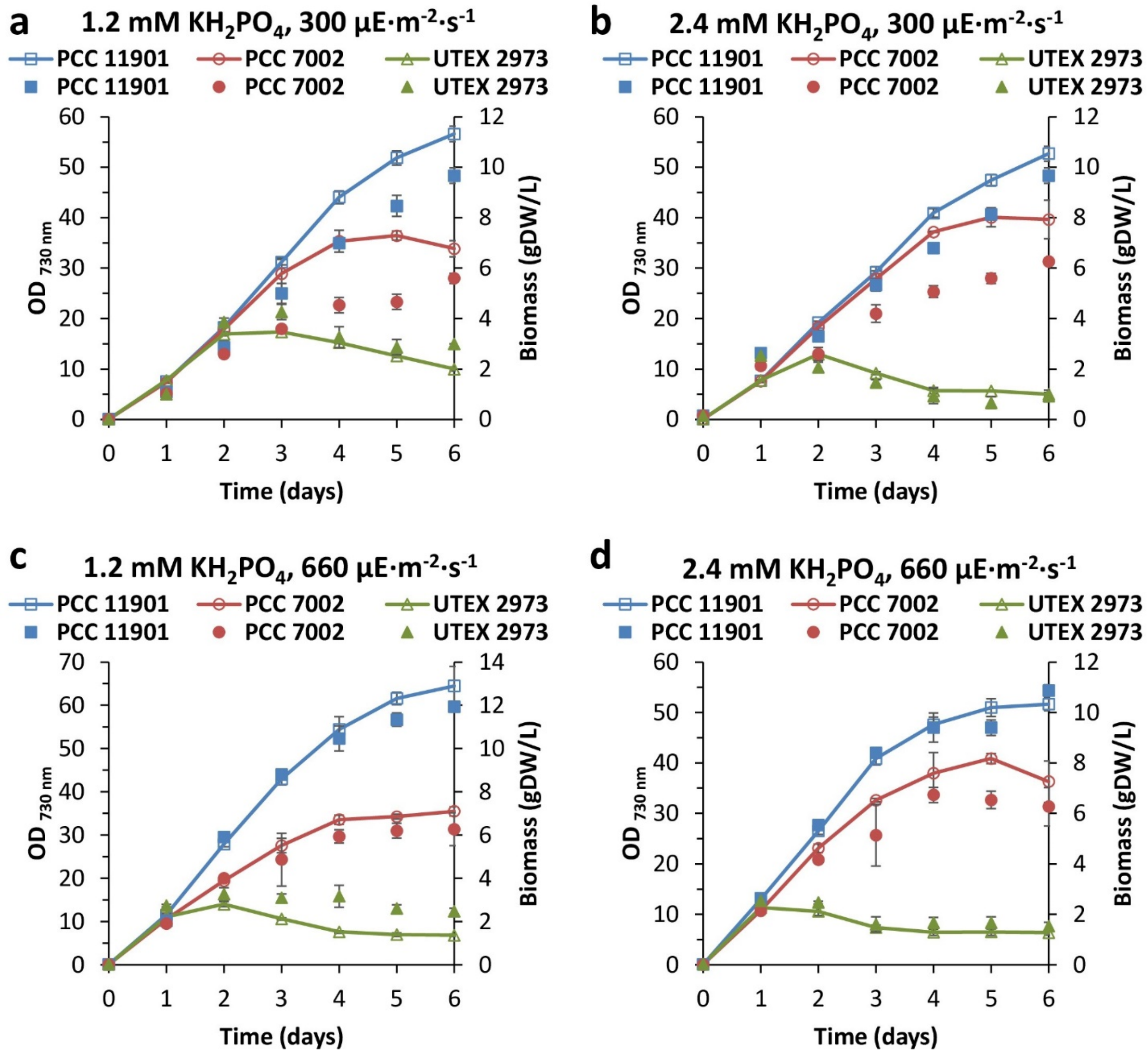


**Figure S6.** Growth performance and biomass accumulation of PCC 11901, PCC 7002 and UTEX 2973 using optimized MAD and MBG media. Lines with empty markers correspond to $OD_{730}$ measurements, whereas filled markers correspond to biomass accumulation. Strains were grown at 38 °C, 225 rpm at constant illumination of either 300 µmol photons·$m^{-2}$·$s^{-1}$ RGB 4:2:1 **(a, b)** or 660 µmol photons·$m^{-2}$·$s^{-1}$ RB 1:1 **(c, d)**. MAD and MBG media used for cultivation of strains contained either 1.2 mM **(a,c)** or 2.4 mM **(b, d)** of $KH_2PO_4$. Average $OD_{730}$ and biomass were calculated as mean of n=3 biological replicates ± standard deviation.

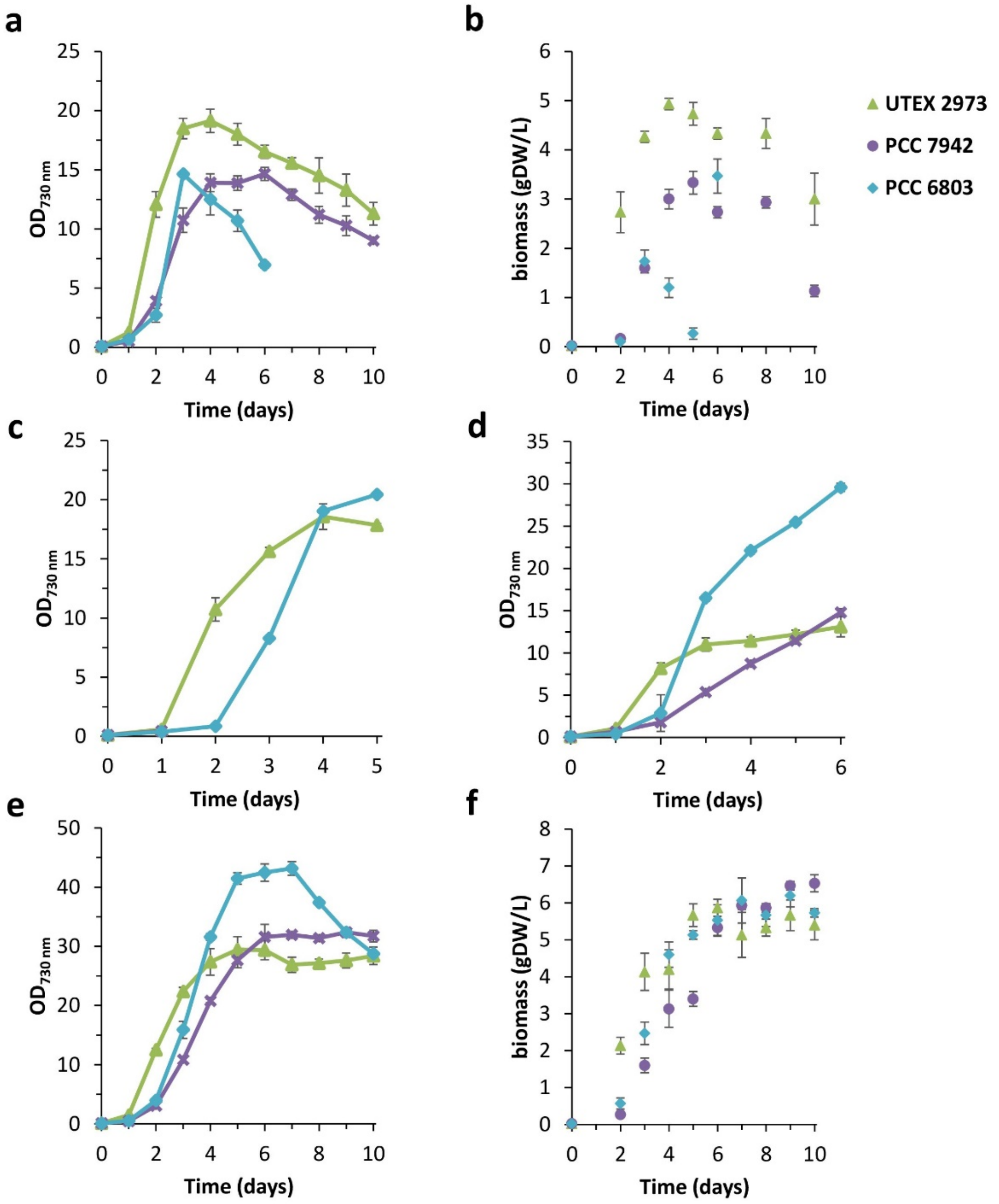


**Figure S7.** Growth and biomass accumulation comparison of UTEX 2973, PCC 7942 and PCC 6803 cultured in different optimized media. All strains were grown at 30 °C and 200 rpm shaking. In the first 24 hours of growth UTEX 2973 was incubated at 150 µmol photons·$m^{-2}$·$s^{-1}$ light intensity, which was then increased to 750 µmol photons·$m^{-2}$·$s^{-1}$. For PCC 7942 and PCC 6803 initial light intensity was set to 75 µmol photons·$m^{-2}$·$s^{-1}$, changed to 150 µmol photons·$m^{-2}$·$s^{-1}$ after 1 day and then increased to 750 µmol photons·$m^{-2}$·$s^{-1}$ on day 2. RGB ratio of LED lights was set to 1:1:1 throughout whole experiment. **(a,b)** Growth and dry cell weight for all freshwater strains grown in MBG medium (BG-11 with 96 mM $NaNO_3$ and 1.2 mM $KH_2PO_4$ and 240 µM ammonium iron (III) citrate) **(c)** MBG supplemented with 20 mM $MgSO_4$, **(d)** MAD medium with 0% NaCl and **(e,f)** 5xBGM (5xBG medium with 96 mM $NaNO_3$, 0.6 mM $KH_2PO_4$ and 0.6 mM $K_2HPO_4$). All above media recipes are listed in **Table S2**. Average $OD_{730}$ and biomass were calculated as mean of n=3 biological replicates ± standard deviation.

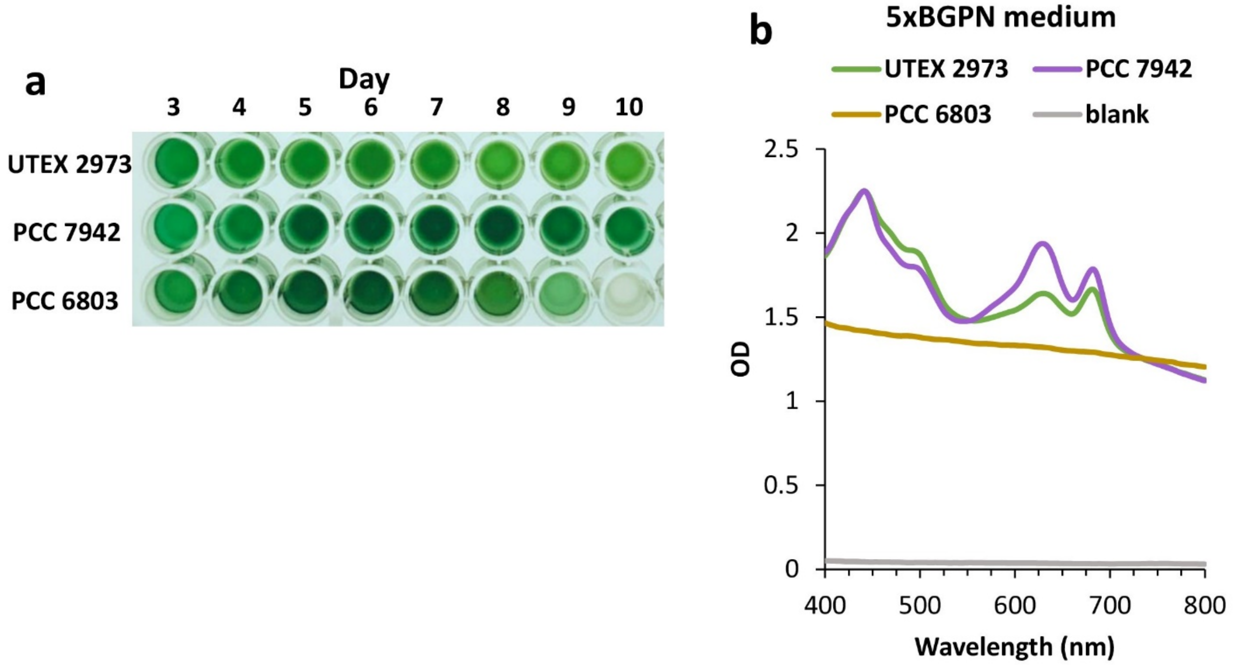


**Figure S8.** Pigmentation of UTEX 2973, PCC 7942 and PCC 6803 cultures in 5xBGM medium. **(a)** Appearance of all strains grown at 30 °C, 200 rpm, 1% $CO_2$ for 10 days. **(b)** Whole cell spectra of cultures collected after 10 days of continuous growth.

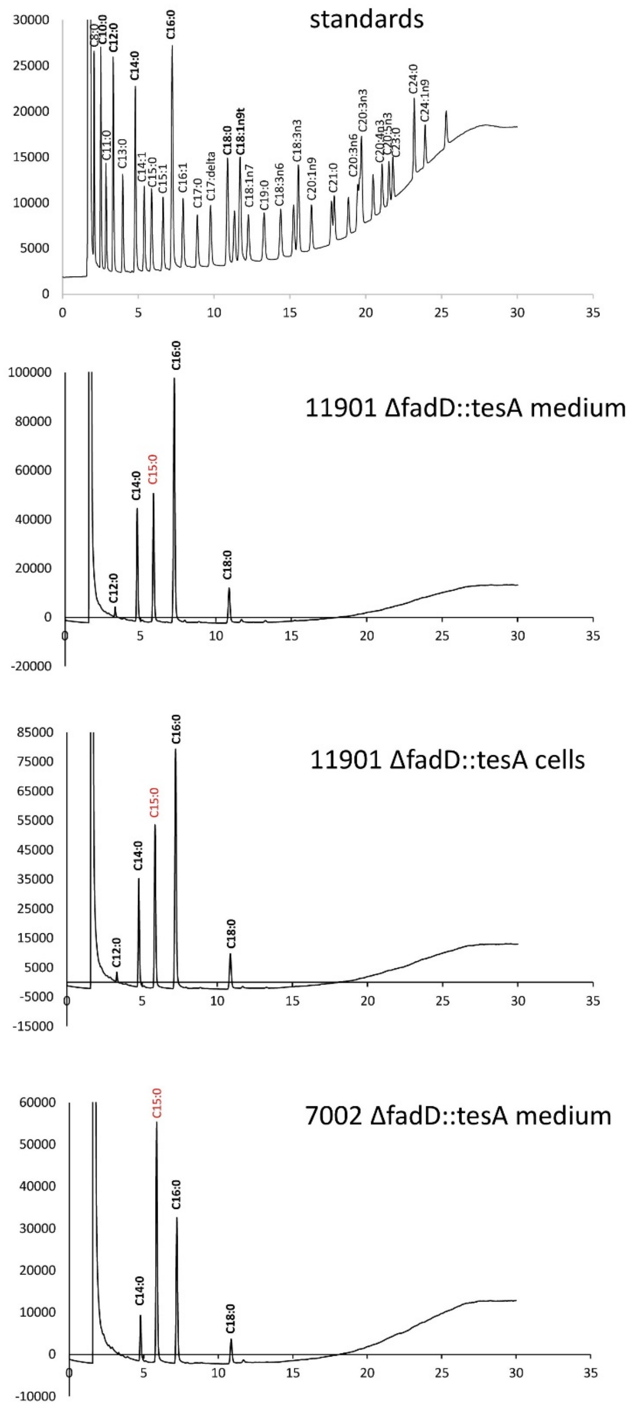


**Figure S9.** GC chromatograms of medium and cell extracts samples of engineered FFA producing strains. Samples were spiked with internal C15 standard.

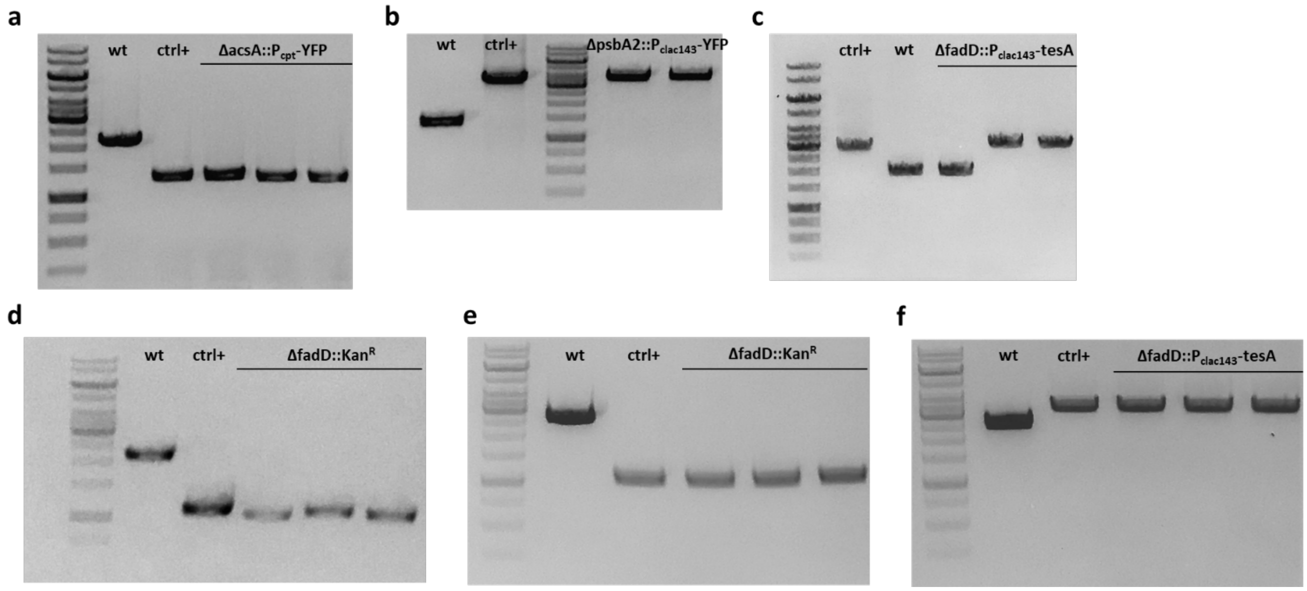


**Figure S10.** Genotyping PCRs of engineered PCC 11901 and PCC 7002 strains. Primers used for the reactions are listed in **Table S4**. Genotyping of **(a)** ΔacsA::$P_{cpt}$-YFP, **(b)** ΔpsbA2::$P_{clac143}$-YFP, **(c)** 11901 ΔfadD::$P_{clac143}$-tesA, **(d)** 11901 ΔfadD::$Kan^R$, **(e)** 7002 ΔfadD::$Kan^R$ and **(f)** 7002 ΔfadD::$P_{clac143}$-tesA transformants. DNA ladder: GeneRuler 1kb (#SM0311, NEB).

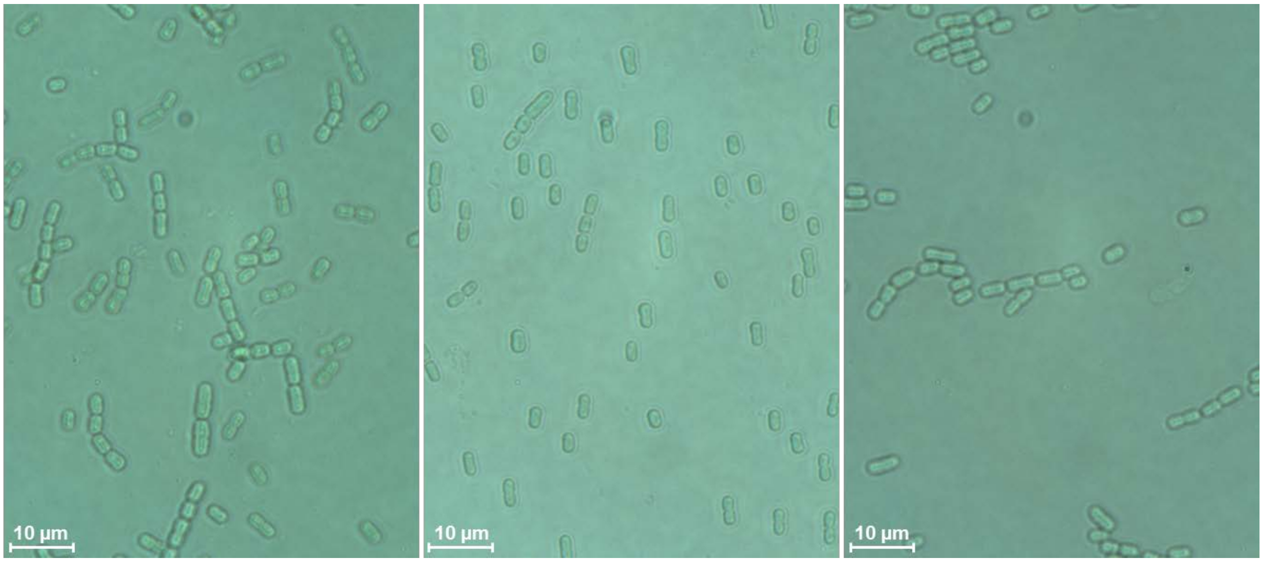


**Figure S11.** Bright-field microscopy images of the PCC 11901 strain (magnification 1000x).

**16S rRNA sequence of the isolated bacterial contaminant:**

GGCAGCTACCATGCAGTCGAGCGCACCTTCGGGTGAGCGGCGGACGGGTTAGTAACGCGTGGGAACGTACCCTT
TTCTAAGGAATAGCCACTGGAAACGGTGAGTAATACCTTATACGCCCTTCGGGGGAAAGATTTATCGGAGAAGGA
TCGGCCCGCGTTAGATTAGATAGTTGGTGGGGTAACGGCCTACCAAGTCTACGATCTATAGCTGGTTTTAGAGGAT
GATCAGCAACACTGGGACTGAGACACGGCCCAGACTCCTACGGGAGGCAGCAGTGGGGAATCTTGGACAATGGG
CGCAAGCCTGATCCAGCCATGCCGCGTGAGTGATGAAGGCCTTAGGGTCGTAAAGCTCTTTCGCTGGGGATGATA
ATGACAGTACCCAGTAAAGAAACCCCGGCTAACTCCGTGCCAGCAGCCGCGGTAATACGGAGGGGGTTAGCGTT
GTTCGGAATTACTGGGCGTAAAGCGCGCGTAGGCGGATTGGAAAGTTGGGGGTGAAATCCCGGGGCTCAACCTC
GGAACTGCCTCCAAAACTATCAGTCTAGAGTTCGAGAGAGGTGAGTGGAATTCCGAGTGTAGAGGTGAAATTCGT
AGATATTCGGAGGAACACCAGTGGCGAAGGCGGCTCACTGGCTCGATACTGACGCTGAGGTGCGAAAGTGTGGG
GAGCAAACAGGATTAGATACCCTGGTAGTCCACACCGTAAACGATGAATGCCAGTCGTCAGCAAGCATGCTTGTT
GGTGACACACCTAACGGATTAAGCATTCCGCCTGGGGAGTACGGTCGCAAGATTAAAACTCAAAGGAATTGACGG
GGGCCCGCACAAGCGGTGGAGCATGTGGTTTAATTCGAAGCAACGCGCAGAACCTTACCAACCCTTGACATCCTG
TGCTACATCCAGAGATGGATGGTTCCCTTCGGGGACGCAGTGACAGGTGCTGCATGGCTGTCGTCAGCTCGTGTC
GTGAGATGTTCGGTTAAGTCCGGCAACGAGCGCAACCCACATCTTCAGTTGCCAGCAGTTCGGCTGGGCACTCTG
GAGAAACTGCCCGTGATAAGCGGGAGGAAGGTGTGGATGACGTCAAGTCCTCATGGCCCTTACGGGTTGGGCTA
CACACGTGCTACAATGGCAGTGACAATGGGTTAATCCCCAAAAACTGTCTCAGTTCGGATTGTTCTCTGCAACTCG
AGAGCATGAAGTCGGAATCGCTAGTAATCGCGTAACAGCATGACGCGGTGAATACGTTCCCGGGCCTTGTACACA
CCGCCCGTCACACCATGGGAGTTGGGTTTACCCGAAGACGGTGCGCCAACCTTTAGGAGGCAGCTGCCACGTANT
ANNNNNNNCCGGCTAACTCCGTGCCAGCAGCCGCGGTAATACGGAGGGGGTTAGCGTTGTTCGGAATTACTG
GGCGTAAAGCGCGCGTAGGCGGATTGGAAAGTTGGGGGTGAAATCCCGGGGCTCAACCTCGGAACTGCCTCCAA
AACTATCAGTCTAGAGTTCGAGAGAGGTGAGTGGAATTCCGAGTGTAGAGGTGAAATTCGTAGATATTCGGAGG
AACACCAGTGGCGAAGGCGGCTCACTGGCTCGATACTGACGCTGAGGTGCGAAAGTGTGGGGAGCAAACAGGAT
TAGATACCCTGGTAGTCCACACCGTAAACGATGAATGCCAGTCGTCAGCAAGCATGCTTGTTGGTGACACACCTAA
CGGATTAAGCATTCCGCCTGGGGAGTACGGTCGCAAGATTAAAACTCAAAGGAATTGACGGGGGCCCGCACAAG
CGGTGGAGCATGTGGTTTAATTCGAAGCAACGCGCAGAACCTTACCAACCCTTGACATCCTGTGCTACATCCAGAG
ATGGATGGTTCCCTTCGGGGACGCAGTGACAGGTGCTGCATGGCTGTCGTCAGCTCGTGTCGTGAGATGTTCGGT
TAAGTCCGGCAACGAACGCAACCCACATCTTCAGTTGCCAGCAGTTCGGCTGGGCACTCTGGAGAAACTGCCCGT
GATAAACCGGAAGGAAGGTGTTGAATGACGTCAAGTCCTCCAGGGCCCTTACCGGGTTGGGGTTACCACCGTGGC
TACAATGGCCAGTGACAATGGGGTTAATCCCCCAAAAACTGTTCCCAGTTCCGAATTGTTTCCTTGCAACTCGGAG
AGCTTGAAATCCGGAA
